# Patient-associated mutations in *Drosophila* Alk perturb neuronal differentiation and promote survival

**DOI:** 10.1101/2022.03.30.486387

**Authors:** K. Pfeifer, G. Wolfstetter, V. Anthonydhason, T. Masudi, B. Arefin, M. Bemark, P. Mendoza-Garcia, R.H. Palmer

**Affiliations:** Department of Medical Biochemistry and Cell Biology, Institute of Biomedicine, Sahlgrenska Academy, University of Gothenburg, SE-405 30 Gothenburg, Sweden; Mucosal Immunobiology and Vaccine Center, Department of Microbiology and Immunology, Institute of Biomedicine, University of Gothenburg, SE-405 30 Gothenburg, Sweden

**Keywords:** neuroblast, tumor, brain, visceral mesoderm, signaling, RTK, neurogenesis

## Abstract

Activating Anaplastic Lymphoma Kinase (ALK) receptor tyrosine kinase (RTK) mutations occur in pediatric neuroblastoma and are associated with poor prognosis. To study ALK-activating mutations in a genetically controllable system we employed CRIPSR/Cas9, incorporating orthologues of the human oncogenic mutations *ALK^F1174L^* and *ALK^Y1278S^* in the *Drosophila Alk* locus. *Alk^F1251L^* and *Alk^Y1355S^* mutant *Drosophila* exhibit enhanced Alk signaling phenotypes, but unexpectedly depend on the Jelly belly (Jeb) ligand for activation. Both *Alk^F1251L^* and *Alk^Y1355S^* mutant larval brains display hyperplasia, represented by increased numbers of Alk-positive neurons. Despite this hyperplasic phenotype, no brain tumors were observed in mutant animals. We show that hyperplasia in *Alk* mutants was not caused by significantly increased rates of proliferation, but rather by decreased levels of apoptosis in the larval brain. Using single-cell RNA sequencing (scRNA-seq), we identify perturbations during temporal fate specification in *Alk^Y1355S^* mutant mushroom body lineages. These findings shed light on the role of Alk in neurodevelopmental processes and highlight the potential of activating Alk mutations to perturb specification and promote survival in neuronal lineages.

## Introduction

Neuroblastoma is the most common and deadly extra cranial solid tumor in children (Maris, 2010; Tolbert & Matthay, 2018). Approximately 10% of neuroblastoma harbor mutations in the Anaplastic Lymphoma Kinase (ALK) receptor tyrosine kinase, providing an important therapeutic target (Trigg & Turner, 2018). Understanding the mechanisms underlying the contribution of ALK mutations to human neuroblastoma development is important, and highly controlled genetic models to investigate their function *in vivo* provide critical insight. Given the ease and control of genetic modification in *Drosophila*, together with the structural conservation of the Alk RTK and the existence of well characterized readouts for Alk activity, we decided to use the fruit fly as model system to interrogate orthologous Alk mutations representing human ALK variants associated with neuroblastoma.

ALK is mainly expressed in the central and peripheral nervous system (Iwahara *et al*, 1997; Vernersson *et al*, 2006). *Alk* knock-out mice are viable and fertile and display mild phenotypes (Bilsland *et al*, 2008; Borenas *et al*, 2021a; Orthofer *et al*, 2020; Rohrbough *et al*, 2013; Witek *et al*, 2015). Knock-in mice harboring neuroblastoma-associated point mutations of *ALK* do not exhibit spontaneous tumor formation after birth but show neuronal hyperplasia (Berry *et al*, 2012; Borenas *et al*, 2021b; Cazes *et al*, 2014; Ono *et al*, 2019). In zebrafish and chicken models, Alk family RTKs are expressed in neural tissues, and have been reported in play a role development of the neural crest (Fadeev *et al*, 2018; Mo *et al*, 2017a; Vieceli & Bronner, 2018; Yao *et al*, 2013) Also, in invertebrate model organisms, such as the fruit fly *Drosophila* and the nematode *C. elegans,* Alk/SCD-2 has roles during neurogenesis (Bazigou *et al*, 2007; Cheng *et al*, 2011; Gouzi *et al*, 2011; Pecot *et al*, 2014). In spite of this, the exact role of ALK in the nervous system and how it relates to development of neuroblastoma is not well understood.

In *Drosophila*, Alk has an indispensable role during embryonic development of the visceral muscles that makes this tissue an excellent model to study Alk function (Englund *et al*, 2003; Lee *et al*, 2003; Stute *et al*, 2004). Alk signaling in response to the Jeb ligand drives specification of founder cell fate, which is critical for visceral muscle fusion (Englund *et al*., 2003; Lee *et al*., 2003; Stute *et al*., 2004). Alk activity in the developing VM is reflected by spatially and temporally regulated phospho-ERK that can easily be detected by antibody staining, providing an ideal readout with which to characterize Alk mutations (Loren *et al*, 2003). This well-characterized Alk-driven embryonic readout can be complemented with analysis of Alk function in the nervous systems at later stages of development. During *Drosophila* larval stages, Alk is robustly expressed in the CNS where it performs multiple functions, including regulation of neuronal targeting and survival, synapse development and body size regulation, brain sparing, longevity, memory, circadian rhythm, ethanol response and learning (Bazigou *et al*., 2007; Cheng *et al*., 2011; Gouzi *et al*, 2018; Gouzi *et al*., 2011; Kumar *et al*, 2021; Lasek *et al*, 2011; Pecot *et al*., 2014; Rohrbough & Broadie, 2010; Rohrbough *et al*., 2013; Weiss *et al*, 2017; Weiss *et al*, 2012; Woodling *et al*, 2020).

*Drosophila* brain development is an excellent model for studying the effect of Alk mutations found in pediatric neuroblastoma patients, as this tumor is considered to arise from aberrant development of precursor cells originating from the neural crest (Hoehner *et al*, 1996; Marshall *et al*, 2014). In the fruit fly, the process of neurogenesis, including the role of neuroblasts (NB) and neural fate specification processes is well characterised, with each NB producing a distinct series of neurons (Yu *et al*, 2006). The central brain is derived from approximately 100 NBs per hemisphere (Urbach & Technau, 2004), that are classified as Type I (NB I) and Type II (NB II). NBs divide asymmetrically to produce different sets of proliferating progeny, with Type I NBs producing ganglion mother cells (GMCs) which subsequently divide into young post mitotic neurons; while Type II NBs generate a larger lineage of neurons via proliferative intermediate precursor cells which divide to generate GMCs that will ultimately divide into young post mitotic neurons (Doe, 2008; Knoblich, 2008; Urbach & Technau, 2004). Diversity of neurons in the *Drosophila* nervous system is achieved by highly regulated consecutive expression of transcription factors, that generate unique combinatorial transcription factor codes within NBs and their progeny (Liu *et al*, 2015; Syed *et al*, 2017a, b; Yang *et al*, 2016). The final complement of neurons is achieved through a binary, Notch dependent manner via apoptosis that occurs after the division of the GMC (Pinto-Teixeira *et al*, 2016).

Alk has previously been described to function within the mushroom body (MB) where it regulates sleep behavior and long term memory formation (Bai & Sehgal, 2015; Gouzi *et al*., 2011). The MB is a bilateral structure containing neurons which arises from four bilateral NBs per brain hemisphere, ultimately producing around 2000 so called Kenyon cells in the adult fly (Crittenden *et al*, 1998). It contains mainly three neuronal cell types that arise in a sequential order during larval and pupal stages: early born γ neurons, middle born α’β’ neurons and late born αβ neurons (Ito *et al*, 1997; Lee *et al*, 1999) and plays distinct roles in olfactory learning, memory and sleep (Cognigni *et al*, 2018; Ito *et al*., 1997; Lee *et al*., 1999). The perikarya of the MB neurons are located on the dorsoposterior surface of the brain and each cell projects their neurites ventroanteriorly into the calyx, formed by the dendrites of the MB neuronal cell bodies. The axons are bundled into the peduncle which branches anteriorly into 5 lobes: two dorsal lobes α and α’, and three medial lobes β, β’ and γ (Crittenden *et al*., 1998). The γ lobe undergoes remodeling during pupation in which its bifurcated structure is pruned due to ecdysone receptor B1 (EcR-B1) activity (Lee *et al*, 2000), whereas the α’ and β’ lobe neurons are specified by expression of the Maternal gene required for mitosis (Mamo) BTB/POZ containing, C2H2 zinc finger transcription factor, which also maintains their cell fate (Liu *et al*, 2019).

Model organisms, such as *Drosophila*, offer a unique opportunity to study the effect of Alk mutations identified in neuroblastoma patients on the process of neuronal fate specification *in vivo*. In pediatric neuroblastoma, failure of differentiation in the neural crest lineage during development of the sympathetic ganglia and the adrenal medulla is thought to result in aggressive neuroblastoma tumors with poor prognosis (Marshall *et al*., 2014; Tomolonis *et al*, 2018). Such tumors often harbor genetic aberrations such as *MYCN* amplification as well as amplification and/or gain-of-function mutations in the ALK RTK (Caren *et al*, 2008; Chen *et al*, 2008; George *et al*, 2008; Hallberg & Palmer, 2013; Janoueix-Lerosey *et al*, 2008; Mosse *et al*, 2008).

Here, we exploit a highly controlled model system, namely *Drosophila melanogaster*, to investigate the effect of *Alk* mutation during neurogenesis. To do this we generated two *Alk* alleles, *Alk^F1251L^* and *Alk^Y1355S^*, which are orthologous to the human *ALK^F1174L^* and *ALK^Y1278S^* mutations in neuroblastoma patients. Remarkably, while the *Alk^Y1355S^* mutant receptor exhibits ectopic Alk signaling in the embryonic visceral mesoderm, this mutant Alk receptor is still ligand dependent. Brains of both *Alk^F1251L^* and *Alk^Y1355S^* exhibit reduced levels of apoptosis resulting in a mild hyperplasia and an environment that is sensitized for tumor formation. However, in agreement with vertebrate models (Borenas *et al*., 2021a; Cazes *et al*., 2014; Ono *et al*., 2019), mutation of *Alk* alone was not sufficient to drive spontaneous tumor development in the *Drosophila* brain. Single cell analysis of *Alk^Y1355S^* larval brains led the identification of differentially expressed genes of which one is Mamo. Since both Mamo and Alk have described functions in the MB (Gouzi *et al*., 2011; Liu *et al*., 2019; Rossi & Desplan, 2020), we focused on Mamo in the MB lineage. Aberrant *Alk^Y1355S^* signaling in the MB lineage leads to precocious Mamo expression in γ neurons during wL3 stage, reflecting defects in neuronal fate specification in mushroom body NB lineages that persist into adulthood. These results provide novel insights into the effect of oncogenic Alk mutations on neuronal specification and survival during development.

## Results

### *Alk^F1251L^* and *Alk^Y1355S^* mutant alleles exhibit gain of function activity

Aberrant signaling of human ALK neuroblastoma variants has previously been characterized by transgenic overexpression in *Drosophila* eye discs, employing a “rough eye” phenotype as readout for ALK receptor activity (Chand *et al*, 2013; Guan *et al*, 2015; Guan *et al*, 2017; Wolfstetter *et al*, 2020). In addition to the rough eye phenotype, the reduced pupal size phenotype observed on overexpression of either *Drosophila* Alk or Jeb ligand in the central nervous system (CNS) also provides a sensitive readout for Alk activity (Gouzi *et al*., 2011; Mendoza-Garcia *et al*, 2017; Wolfstetter *et al*, 2017). To address whether the ability of oncogenic human ALK to drive the reduced pupal size phenotype is conserved, we employed the pan neuronal *C155-Gal4* driver to overexpress either (i) wild type human ALK, (ii) human ALK-F1174L gain-of-function (representing one of the three ALK hotspot mutations in neuroblastoma) or (iii) an additional ALK-Y1278S gain-of-function neuroblastoma mutant (Umapathy *et al*, 2019). Expression of either ALK-F1174L or ALK-Y1278S activated mutant variants, but not wild type ALK, resulted in a reduced pupal size phenotype, suggesting that ALK receptor signaling output is conserved between human and *Drosophila* (Fig. 1A). However, while informative, overexpression results in substantial modification of receptor numbers and molecular signaling dynamics that can potentially lead to non-specific phenotypes. We therefore decided to generate mutations in the endogenous *Alk* locus that model neuroblastoma patient point mutations in a controlled genetic background, and two representative mutations – human *ALK-F1174L* and *ALK-Y1278* – were selected for this purpose. These gain-of-function ALK kinase mutants are reported as constitutively active ALK mutations found in human neuroblastoma patients, with *ALK-F1174L* being a more frequently occurring “hot-spot” mutation (Fig.1B-C) (Umapathy *et al*., 2019). Sequence alignment analysis identified the equivalent residues in *Drosophila* Alk as *Alk-F1251* and *Alk-Y1355* (Fig. 1B-C). A CRISPR/Cas9 mediated homologous donor repair (HDR) strategy was used to generate *Alk^F1251L^* (phenylalanine to leucine at amino acid position 1251) and *Alk^Y1355S^* (tyrosine to serine at position 1355) mutations in the *Drosophila Alk* locus that would allow us to model the effect of these patient derived gain-of-function human ALK neuroblastoma mutations in the fly brain.

**Figure 1.**
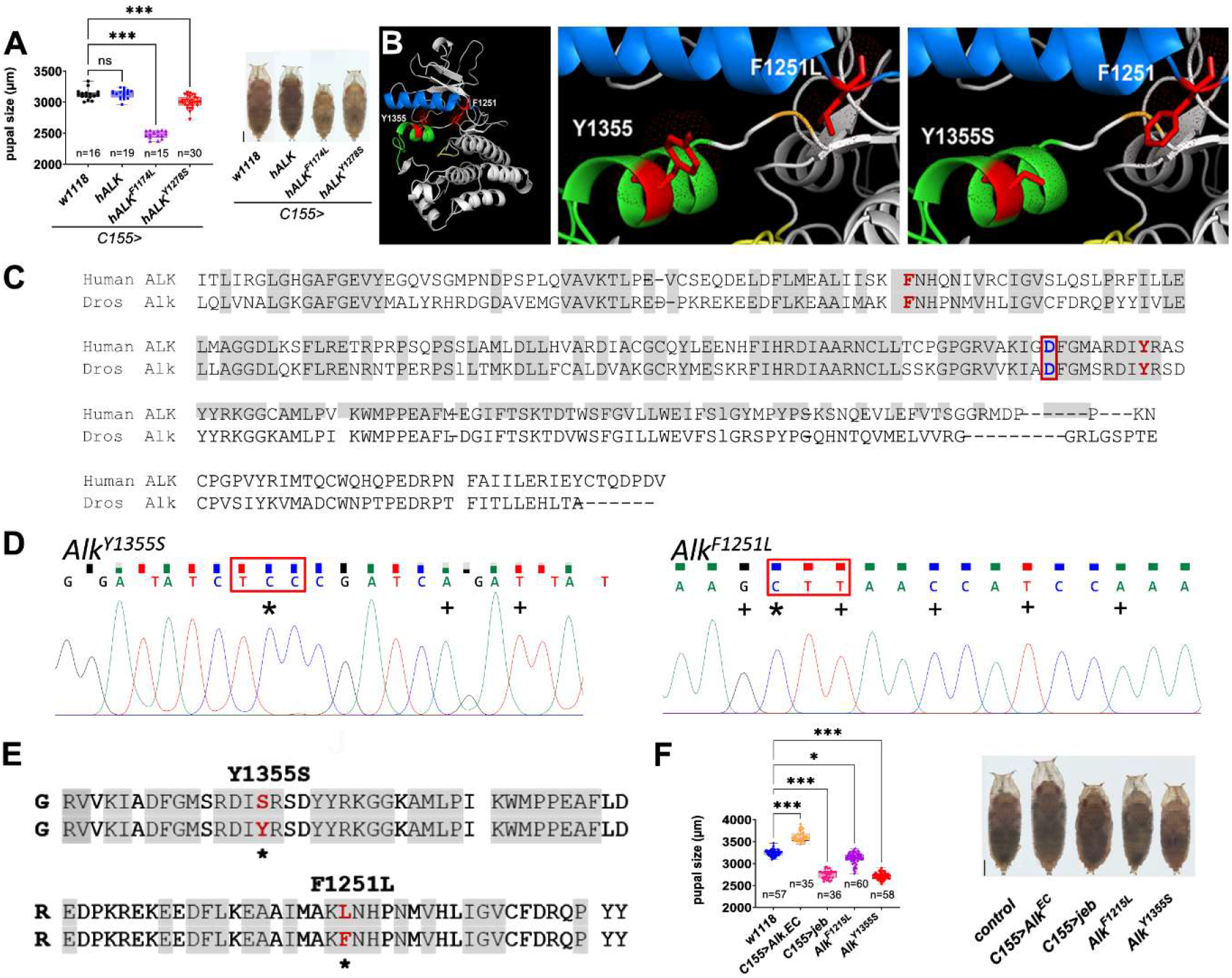
CRISPR/Cas9-mediated generation of neuroblastoma-associated *Alk* mutants in *Drosophila*. **A** Pupal size analysis. Expression of human ALK gain-of-function transgenes (*C155>hALK^F1174L^* or *C155>hALK^Y1278S^*), but not wild type human ALK (*C155>hALK*) in the *Drosophila* CNS leads to a significant change in pupal case size. Representative examples are shown. Statistics: One-way-ANOVA, Dunnett’s multi comparison test; *** = *p*<0.001. Scale bar 500 µm. **B** Structural model of the *Drosophila* Alk kinase domain indicating residues F1251 and Y1355. In *blue*: alpha-C helix, *green*: activation loop *red*: amino acids F1251 and Y1355. **C** Protein sequence alignment of human and *Drosophila* Alk in the kinase domain, with conserved regions highlighted in grey. Two mutated residues found in human neuroblastoma, F1174 and Y1278 (orthologous to *Drosophila* F1251 and Y1355, respectively), are indicated in red. The D1268 (orthologous to *Drosophila* D1345) residue predicted to be essential for kinase activity is indicated in blue with a red box. **D** Chromatogram confirming CRISPR/Cas9-mediated nucleotide exchange (asterisks) in the *Alk^F1251L^* and *Alk^Y1355S^* mutant alleles, + indicates silent mutations. **E** Amino acid alignment of the generated *Alk^F1251L^* and *Alk^Y1355S^* mutants. Upper sequence is wild type. Asterisk marks amino acid changes. **F** Pupal size analysis. Activation (*Alk^F1251L^*, *Alk^Y1355S^, C155>jeb*) or inhibition (*C155>Alk.EC*) of Alk signaling in the CNS leads to a significant change in pupal case size. Representative examples are shown. Statistics: One-way-ANOVA, Kruskal-Wallis test; *** = *p*<0.001. Scale bar 500 µm.

Both mutations were successfully generated in the kinase domain encoding portion of the *Alk* locus and verified by sequencing (Fig. 1D-E). Flies harbouring *Alk^F1251L^* and *Alk^Y1355S^* alleles were viable both as heterozygous and homozygous stocks and were subsequently characterized for gain-of-function Alk activity in two independent assays: (i) analysis of pupal size and (ii) analysis of Alk signaling in the embryonic visceral mesoderm. As previously reported, activation of Alk by Jeb ligand expression in the nervous system (*C155>jeb*) resulted in a reduced pupal size phenotype (Fig. 1F), while inhibition of Alk signaling by overexpression of a dominant negative Alk variant (*C155>Alk.EC*) increased pupal size (Fig. 1F). Both *Alk^F1251L^* and *Alk^Y1355S^* alleles displayed a reduced size phenotype indicating that both mutations are indeed Alk gain-of-function mutations (Fig. 1F). Interestingly, we note that (i) *Alk^Y1355S^* shows a stronger phenotype than *Alk^F1251L^*, indicating that these mutations are not equally strong gain-of-function mutations and (ii) heterozygous (e.g. *Alk^Y1355S^/+*) pupae display a weaker phenotype than homozygous mutants, suggesting dosage sensitivity (Suppl. Fig. 1A).

To further analyze Alk signaling in the *Alk^F1251L^* and *Alk^Y1355S^* mutant alleles we turned to the embryonic visceral mesoderm (VM), where activation of Alk signaling by Jeb leads to the specification of founder cells which can be visualized by phospho-ERK staining (Englund *et al*., 2003; Lee *et al*., 2003; Loren *et al*., 2003; Stute *et al*., 2004) or *HandC-GFP* reporter expression (Fig. 2A-A’,F,F’). On Jeb overexpression, all Alk expressing cells in the VM respond with robust phospho-ERK staining and *HandC-GFP* reporter activation (Fig. 2B and G). In contrast, no detectable phospho-ERK staining or *HandC-GFP* reporter expression was observed when Alk signaling is perturbed in an *Alk* kinase domain deletion mutant (Wolfstetter *et al*., 2017) (*Alk^KO^*, Fig. 2C and H). Analysis of Alk signaling in the VM, employing both phospho-ERK and the *HandC-GFP* reporter as readouts for Alk activation, shows that the *Alk^Y1355S^* mutation results in ectopic Alk signaling (Fig. 2D and I) in contrast to the restricted founder cell activation observed in wild type (Fig. 2A-A’, F-F’). Surprisingly, no difference was seen between the Alk signaling output in the VM of *Alk^F1251L^* mutants and control embryos (Fig. 2E and J). Taken together, these findings suggest that the *Alk^F1251L^* and *Alk^Y1355S^* alleles generated here represent *Drosophila Alk* gain-of-function alleles of differing strength, with *Alk^Y1355S^* representing a stronger gain-of-function allele than *Alk^F1251L^*.

**Figure 2.**
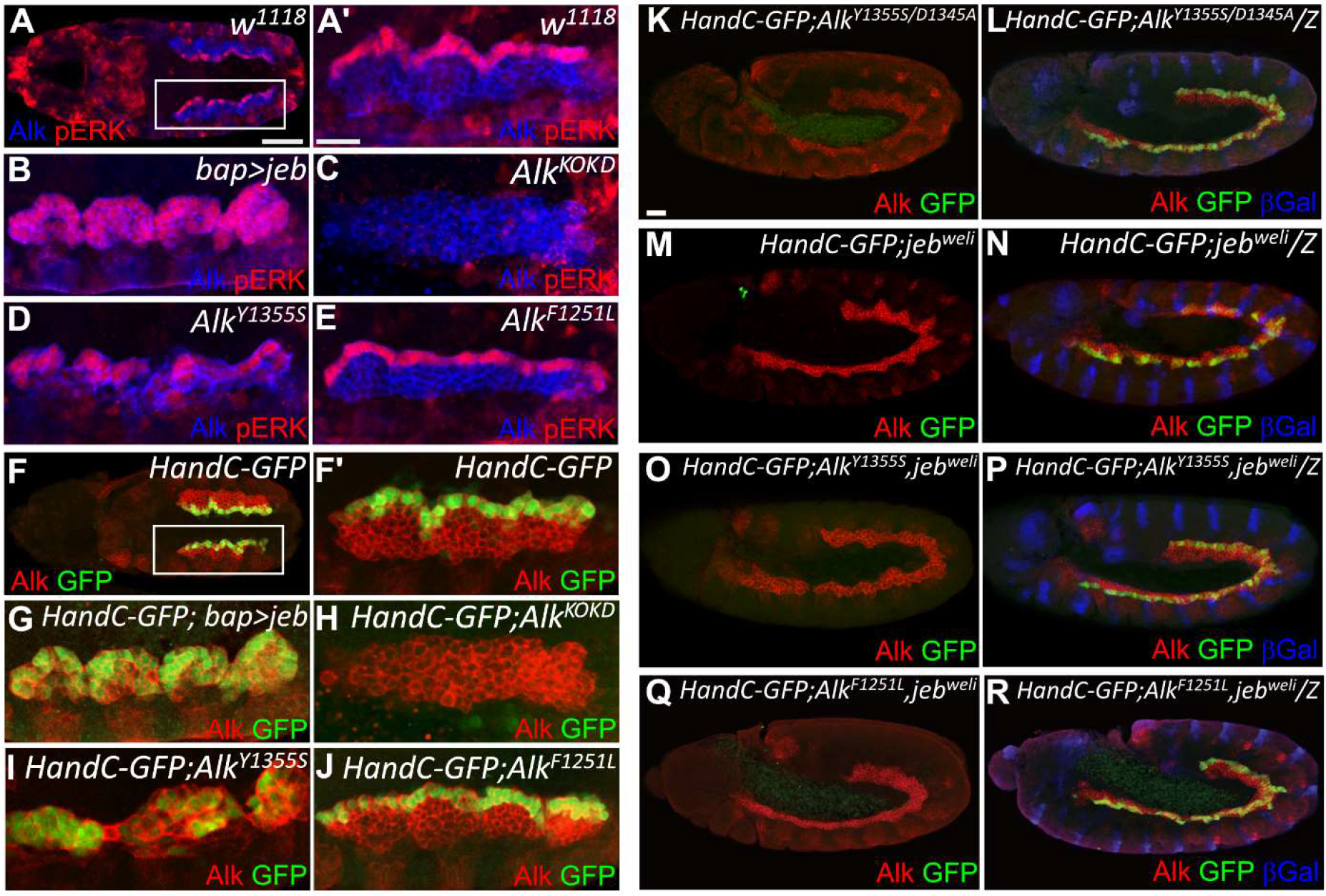
Analysis of Alk signaling in the embryonic visceral mesoderm of *Alk^F1251L^* and *Alk^Y1355S^* mutant alleles. **A-A’** Visualisation of Alk signaling in wild type embryonic founder cells (FCs, dorsal row) of the visceral mesoderm (VM) with anti-phospho-ERK (A’ - magnification of boxed region in A). Scale bars 50 µm (A) and 10 µm (A’). **B** Overexpression of Jeb with *bap-Gal4* leads to Alk signaling within the whole VM. **C**. No phospho-ERK is observed in an *Alk* loss-of-function mutant (*Alk^KO^*; (Wolfstetter *et al*., 2017)). **D** The *Alk^Y1355S^* mutant allele displays increased phospho-ERK in VM cells. **E** No detectable change in phospho-ERK staining was observed in *Alk^F1251L^*. **F-J** Similar results are observed when the *HandC-GFP* reporter is employed as readout for Alk signaling in the VM. **F-F’** Wild type embryos display robust expression of *HandC-GFP* in VM FCs. F’ magnification of F. **G-H** Overexpression of Jeb results in *HandC-GFP* expression in all VM cells (G), while *HandC-GFP* expression is absent in *Alk^KO^* (H). **I** Ectopic *HandC-GFP* positive VM cells are observed in *Alk^Y1355S^*. **J** *HandC-GFP* expression in *Alk^F1251L^* is similar to controls. **K** *Alk^Y1355S, D1345A^* homozygous double mutants do not show *HandC-GFP* reporter expression in the VM. Scale bar 20 µm. **L** Expression of *HandC-GFP* in *Alk^Y1355S, D1345A^*/*CyO, wg-lacZ* VM FCs. **M-N** *HandC-GFP* reporter expression in *jeb^weli^* homozygous mutant embryos and heterozygous control embryos at stage 11. **O, Q** *HandC-GFP* expression in *Alk^Y1355S^*, *jeb^weli^* and *Alk^F1251L^, jeb^weli^* homozygous mutant embryos at stage 11. **P, R** *Alk^Y1355S^*, *jeb^weli^* and *Alk^F1251L^, jeb^weli^* heterozygous controls at stage 11.

### *Alk^Y1355S^* and *Alk^F1251L^* mutants are ligand dependent

To ensure the phenotypes observed were specific for modification of the *Alk* locus, we combined a kinase-dead mutation (*Alk^D1345A^*) with the *Alk^Y1355S^* allele using CRISPR/Cas9-mediated HDR. *Alk^D1345A^* harbors a modification of the highly conserved DFG motif in the kinase domain, in which the ATP-gamma-phosphate-binding aspartic acid 1345 is changed to alanine (Fig. 1C). As expected, homozygous *Alk^D1345A^* animals were lethal and failed to specify embryonic VM founder cells (Suppl. Fig.1B). Analysis of the *Alk^D1345A, Y1355S^* double mutant allele showed that the embryonic VM phenotype of *Alk^Y1355S^* was abrogated by the D1345A kinase dead mutation, resulting in an *Alk* loss-of-function phenotype (Fig. 2K-L). Further, although heterozygous *Alk^D1345A^/+* pupae did not exhibit a size phenotype, the reduced size phenotype of *Alk^Y1355S^/+* was rescued in *Alk^D1345A, Y1355S^/+*animals (Suppl. Fig.1A). Taken together, these data confirm that (i) Alk kinase activity is required for signaling output in both wild-type and gain-of-function backgrounds, and that (ii) the phenotypes observed in the *Alk^Y1355S^* mutant allele are specific effects of increased Alk signaling and not due to CRISPR/Cas9 off target effects.

We next investigated whether the *Alk^F1251F^* and *Alk^Y1355S^* alleles were ligand independent by testing their ability to rescue *jeb* loss-of-function mutants, which also fail to develop the visceral musculature leading to an non-functional midgut (Englund *et al*., 2003; Lee *et al*., 2003; Stute *et al*., 2004; Weiss *et al*, 2001) (Fig. 2M compared to control Fig. 2N). Remarkably, *Alk^Y1355S^, jeb^weli^* double mutants were lethal and exhibited the *jeb^weli^* phenotype (Fig. 2O compared to control Fig. 2P). We also tested the ability of *Alk^F1251L^* to rescue *jeb* mutants, confirming that this mutant allele is also ligand dependent (Fig. 2Q-R). These data clearly show that while both *Alk^Y1355S^* and *Alk^F1251L^* mutant Alk RTKs exhibit gain-of-function activity in terms of pupal case size they remain ligand dependent in the context of embryonic VM founder cell specification.

### Alk is strongly expressed in mature neurons of the *Drosophila* larval brain

Previous studies have reported broad expression of Alk mRNA and protein in the *Drosophila* nervous system (Bazigou *et al*., 2007; Cheng *et al*., 2011; Gouzi *et al*., 2011; Loren *et al*., 2003; Loren *et al*, 2001). We used fluorescent *in situ* hybridization chain reaction (HCR FISH) to detect *Alk* and *jeb* mRNA in the CNS of control (*w^1118^*) and *Alk^Y1355S^* animals. Both *Alk* and *jeb* mRNA were robustly expressed at similar levels in both control and *Alk^Y1355S^* third instar larval brains (Fig. 3A). To further investigate the effect of the *Alk^Y1355S^* mutation, we analyzed third instar larval brains of control (*w^1118^*) and *Alk^Y1355S^* alleles using 10X genomic based single-cell RNA sequencing (scRNA-seq). In total, scRNA-seq data was collected for 3967 control and 4099 *Alk^Y1355S^* high quality CNS cells and further analyzed with R (Seurat) and Python-based (Scanpy) pipelines. This analysis initially defined 19 cell clusters (Suppl. Fig. 2 A), which were merged based on canonical markers (Suppl. Fig. 2B and Suppl. Fig. 2D), resulting in 8 distinct clusters on a UMAP based 2D projection (Fig. 3B). These 8 clusters were defined as: Early Neuroblast, Neuroblast Enriched Cells, Immature Neurons, Mature Neurons, Neuroblast Proliferating Cells, OLE, Repo +ve cells and Wrapper +ve cells. Canonical markers for these subsets were expressed in these clusters (Fig. 3C and Suppl. Fig. 2D), and we identified several other unique marker genes for each subtype (Ariss *et al*, 2020; Brunet Avalos *et al*, 2019; Cattenoz *et al*, 2016; Estacio-Gomez *et al*, 2020; Michki *et al*, 2021)(Suppl. Fig. 2E). In addition to these clusters we identified two additional clusters representing hemocytes (cluster 3 in Suppl. Fig. 2A) and cuticle associated cells (cluster 18 in Suppl. Fig. 2A), defined by the *He*, *Idgf6*, *Tep4*, *Hml*, *dpy*, *Obst-B*, *Cht10* and *pot* markers, respectively (Suppl. Fig. 2C) (Cattenoz *et al*, 2020; Dong *et al*, 2020; Evans *et al*, 2014; Öztürk-Çolak *et al*, 2016).

**Figure 3.**
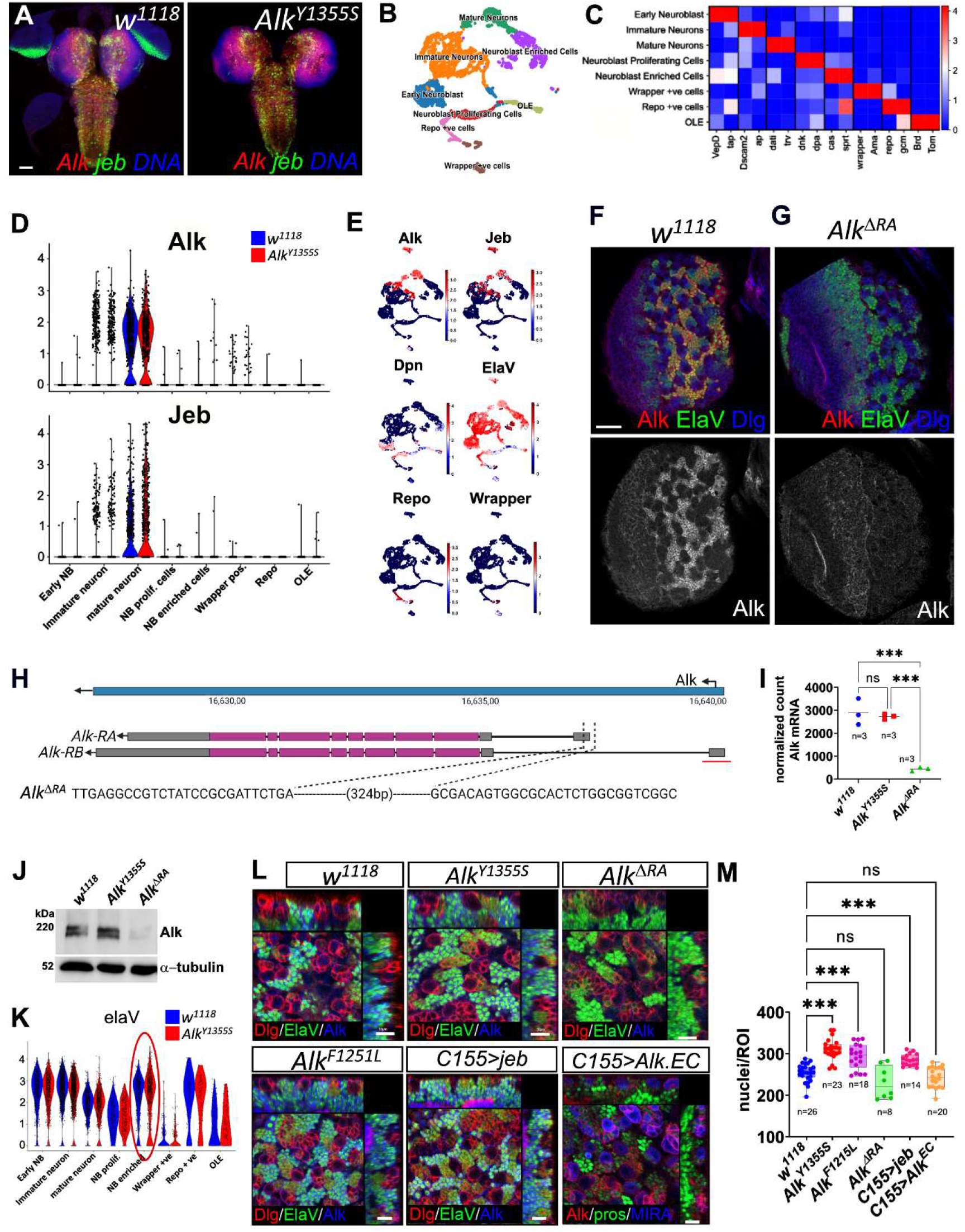
*Alk^F1251L^* and *Alk^Y1355S^* larval brains display hyperplasia. **A** Expression of *Alk* (red) and *jeb* (green) mRNAs *in situ* are shown, DAPI in blue. **B** UMAP displaying the cellular heterogeneity of the wild type scRNA-seq dataset as eight cell types (early neuroblast, neuroblast enriched cells, immature neurons, mature neurons, neuroblast proliferating cells, optic lobe epithelium (OLE), repo-positive cells and wrapper-positive cells). **C** Matrix plot visualizing canonical markers (two per cluster) defining the cellular heterogeneity of the scRNA-seq dataset. Scale bar indicates mean expression values of gene expression. **D** Violin plots showing the mRNA expression of *Alk* and *jeb* across larval brain scRNA-seq clusters (blue denotes wild type and red denotes *Alk^Y1355S^*). **E** Feature plots visualising expression of *Alk, jeb, ElaV, dpn, repo* and *wrapper* within the single cell population. Color scale indicates expression level. **F-G** Alk protein expression in the central brain area (one wL3 brain hemisphere is shown), in *w^1118^* control (F) and *Alk^ΔRA^* mutant (G). Alk (red), ElaV (green) and Dlg (blue) are shown. Scale bar 20 µm. **H** Schematic outlining the genomic organization of the *Alk* locus (blue). Intron-exon structure of both *Alk-RA* and *Alk-RB* transcripts is shown below. Open reading frame in purple. The sequence of the *Alk^ΔRA^* CRISPR/Cas9 deletion mutant is shown below. The region deleted in the previously described *Alk^ΔRB^* CRISPR/Cas9 deletion mutant (Mendoza-Garcia *et al*., 2017) is indicated for reference as red line. **I** *Alk* mRNA expression levels in wild-type, *Alk^Y1355S^* and *Alk^ΔRA^* third instar larval brain bulk RNA-seq data. **J** Western blot of wL3 larval brains indicating Alk protein levels in control (*w^1118^*), *Alk^Y1355S^* and *Alk^ΔRA^* third instar larval brain lysates. Tubulin is employed as loading control. **K** Violin plot indicating *ElaV* mRNA expression across the third instar larval brain scRNA-seq clusters. Red circle highlights expression of *ElaV* in the NB enriched cluster in *Alk^Y1355S^*. **L** Orthogonal projections of the wL3 central brain area showing ElaV-positive perikarya at the NB level. Scale bar 10 µm. Quantification and statistical analysis is shown in **M**, indicating increased numbers of nuclei per region of interest (ROI) in *Alk^Y1355S^* and *C155>jeb* animals. No significant changes in nuclei per ROI were observed in either *Alk^ΔRA^* or *C155>Alk^EC^*. Statistics: One-way Anova, Dunnett’s multiple comparison test; ***p<0.001.

*Alk* and *jeb* were robustly expressed in the scRNA-seq dataset of both control and *Alk^Y1355S^* larval brains. Both *Alk* and *jeb* were predominantly expressed in mature neuronal lineages (Fig. 3D-E), including cholinergic, glutamatergic and GABAergic neurons (Suppl. Fig. 2F). This is in keeping with the previously described strong expression of Alk in the medulla neuropile of the optic lobe (Bazigou *et al*., 2007), and of Jeb in the cholinergic neurons (Okamoto & Nishimura, 2015). Interestingly, we detected very little *Alk* or *jeb* at the mRNA level in either glial cell or neuroblast enriched cell populations (Fig. 3D-E).

We next investigated Alk protein expression in the third instar larval brain, confirming that Alk protein was strongly expressed in ElaV-positive progeny of both Type I and Type II neuroblasts (NBs) (Fig. 3F). To examine the role of Alk further, we generated an *Alk* mutant allele (*Alk^ΔRA^*), specifically removing *Alk* expression in the larval brain (Fig. 3G-H). In previous work, we have shown that the promoter associated with *Alk-RB* isoform expression (red line in Fig. 3H) is essential for expression of Alk in the developing embryonic visceral mesoderm (Mendoza-Garcia *et al*., 2017). Employing isoform specific probes to detect both *Alk-RA* and *Alk-RB* transcripts we confirmed the expression of the previously described *Alk-RB* isoform (Mendoza-Garcia *et al*., 2017) in the *Drosophila* embryonic visceral mesoderm and showed that the *Alk-RA* isoform is expressed in the epidermis and developing nervous system (Suppl. Fig. 3A and B). Based on this we employed CRISPR/Cas9-mediated targeted deletion to remove the *Alk-RA* transcript generating the *Alk^ΔRA^* mutant allele (Fig. 3H). Indeed, expression of Alk protein in the third instar larval central brain was significantly reduced, which was further confirmed by RNA-seq analysis (Fig. 3G and I) and immunoblotting (Fig. 3J). *Alk^ΔRA^* mutants were viable with no obvious defects during development, providing an excellent tool with which to further investigate Alk function in the larval brain.

### *Alk^F1251L^ and Alk^Y1355S^* exhibit neuronal hyperplasia at larval stages

Since our scRNA-seq dataset and subsequent Alk antibody staining highlighted robust expression of Alk in neurons of the central brain, we focused on this area for further investigation. A closer examination of the central brain area of *Alk^Y1355S^* and *Alk^F1251L^* identified increased numbers of neurons relative to controls (Fig. 3L). This increase in ElaV-positive neurons was also observed on pan-neuronal expression of Jeb with *C155-Gal4* (Fig. 3L). Loss of Alk activity, either by overexpression of dominant negative Alk *(C155>Alk.EC*) or in the *Alk^ΔRA^* mutant (Fig. 3L) did not significantly affect the numbers of ElaV-positive neurons. These results were further strengthened by a slight increase in ElaV expressing cells in the *Alk^Y1355S^* scRNA-seq dataset (Fig. 3K). To quantify this phenotype, third instar larval brains were stained with DAPI, and perikarya within a region of interest (ROI) at the neuroblast level counted. A significantly increased number of nuclei were observed in both *Alk^Y1355S^* and *Alk^F1251L^* mutant alleles as well as on Jeb overexpression (*C155>Jeb*) (Fig. 3M). Notably, these observations are in keeping with observations of hyperplasia in the nervous system of *Alk* gain-of-function mouse models (Borenas *et al*., 2021b; Cazes *et al*., 2014). No significant change in number of neurons was observed in third instar larval brains lacking Alk activity, either employing an Alk dominant negative (*C155>Alk.EC*) or the *Alk^ΔRA^* mutant allele, suggesting that Alk function is not critically required for their differentiation. Thus, our data suggests that the larval brains of *Alk* gain-of-function alleles contain increased numbers of mature neurons.

### Single cell RNAseq analysis identifies perturbed neuronal identity in *Alk^Y1355S^* larval brains

Integration of *Alk^Y1355S^* and control scRNA-seq datasets was performed to identify shared and distinct cell identities (Fig. 4A), highlighting an increased number of cells in the “neuroblast enriched” cluster (Fig. 4B), further suggesting that increased Alk activation may result in a perturbation of neuroblast derived lineages and populations. We therefore interrogated this cell cluster in our scRNA-seq datasets further. Re-analysis of the neuroblast enriched cell cluster resulted in 5 cell clusters (Fig. 4C-D). While cluster 1 was enriched in *Alk^Y1355S^* compared to the control scRNA-seq dataset, similar numbers of *Miranda (Mira)*-positive neuroblasts (clusters 3 and 4), were observed and antibodies to Mira confirmed similar numbers of Mira-positive neuroblasts in *Alk^Y1355S^* and control larval brains (Fig. 4E). We also counted NBs per brain hemisphere in *Alk^Y1355S^*, *Alk^F1251L^* and *Alk^ΔRA^*, but were unable to observe any change in overall NB number in *Alk* mutant brains, indicating that the additional neurons observed in *Alk^Y1355S^* and *Alk^F1251L^* brains were not a result of elevated NB numbers (Fig. 4F).

**Figure 4.**
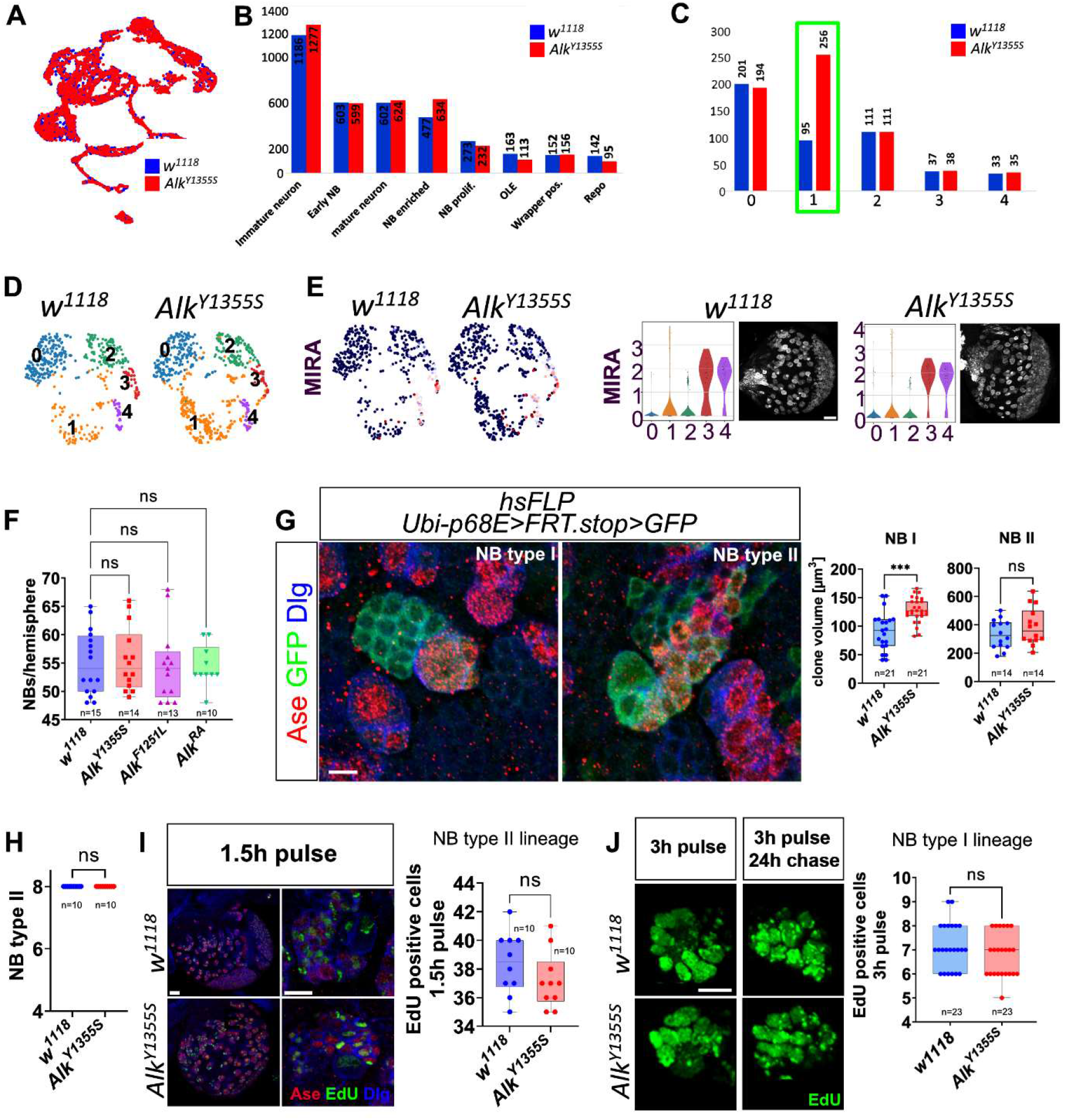
Single cell analysis identifies perturbed neuroblast lineages in *Alk^Y1355S^* mutants. **A** UMAP of the integrated scRNA-seq control (*w^1118^*) (blue) and *Alk^Y1355S^* (red) datasets. **B** Bar chart indicating the number of cells in each cluster across the whole brain cell population in both control (*w^1118^*) (blue) and *Alk^Y1355S^* (red) scRNA-seq datasets. **C** Bar chart displaying cell distribution in a reanalysis of the neuroblast enriched cell population in B. Increased numbers of cells are observed in cluster 1 (green rectangle) of *Alk^Y1355S^* (red), compared to control (blue). **D** UMAPs showing the five sub-clusters (shown in C) identified in control (*w^1118^*) and *Alk^Y1355S^* from the NB enriched population. **E** UMAPs indicating *miranda (Mira)* mRNA expression in both wild-type and *Alk^Y1355S^* scRNA-seq datasets. Violin plots show *Mira* mRNA expression and distribution in the subclusters of the neuroblast enriched cell population of control (*w^1118^*) and *Alk^Y1355S^*, together with Mira expression in wL3 larval brains. Scale bar 20 µm. **F** Quantification of NBs per brain hemisphere in *Alk^Y1355S^*, *Alk^F1251L^* and *Alk^ΔRA^* compared to controls (*w^1118^*). No significant changes in NB numbers were observed. One-way Anova, Turkey’s multiple comparisons test, p=not significant. **G** GFP-positive clones derived from type I NBs in either control (*w^1118^*) or *Alk^Y1355S^* mutant backgrounds. Scale bar 5 µm. Quantification shown at right, n = 10 per genotype. Significant differences in clonal volume were observed in *Alk^Y1355S^*. Statistics: Unpaired t test; ***p<0.001. **H** Total number of Type II NBs per brain hemisphere in control (*w^1118^*) and *Alk^Y1355S^*. Unpaired t test, Kolmogorov-Smirnov test, p=not significant (ns). **I** Quantification of EdU-positive cells in Type II NB lineages as identified by anti-Asense staining for control (*w^1118^*) and *Alk^Y1355S^* wL3 larval brains. No significant differences were observed, Unpaired t test p=not significant (ns). Scale bars 20 (left panel) and 5 µm (right panel). **J** EdU pulse-chase experiments show no significant difference in EdU-positive cells arising from NB type I lineages (n=23) in *Alk^Y1355S^* larval brains compared with control (n=23). EdU incorporation was quantified for 3 h pulse and 3 h pulse plus 24 h chase conditions. Unpaired t test, Mann-Whitney test p>0.2013 (not significant, (ns)). Scale bar 5 µm.

To experimentally address the dynamics of neuroblast lineages, GFP-marked clones were generated in both control (*w^1118^*) and *Alk^Y1355S^* backgrounds (Fig. 4G). Analysis of the clones descended from Type I NBs revealed significant increase in clone volume in *Alk^Y1355S^* (Fig. 4G), in agreement with our earlier observation of increased numbers of ElaV-positive perikarya in *Alk^Y1355S^* larval brains. Analysis of Type II neuroblast clone size showed a non-significant trend towards increased clone volume (Fig. 4G), which was also reflected by similar numbers of Type II NBs per brain lobe (Fig. 4H). We further employed EdU pulse chase analysis, subjecting *Alk^Y1355S^* and control (*w^1118^*) larvae to either (i) EdU pulse (1.5 h for NB II and 3 h for NB I) followed by immediate dissection or (ii) EdU pulse followed by 24 h chase. EdU pulse chase analyses did not reveal any significant differences in proliferation of Type I or Type II NBs when comparing *Alk^Y1355S^* larval brains with *w^1118^* controls (Fig. 4I-J) indicating that the *Alk^Y1355S^* mutation does not increase proliferation rates in larval brains. Moreover, we employed the FUCCI-system in both L3 larvae after eclosion and wandering L3 larvae which enabled us to identify cells in different phases of the cell cycle (Zielke *et al*, 2014) (Suppl. Fig. 4 and 5). We were unable to see differences in cell cycle phases in *Alk^Y1355S^* compared to the control, indicating that proliferation is not enhanced in the mutant at the examined time point. Thus, our single cell and subsequent experimental validations suggest that the hyperplasia observed in *Alk^Y1355S^* larval brains is not due to increased NB numbers or proliferation, implying other mechanisms are responsible.

### Neuroblast quiescence proceeds as normal in *Alk^Y1355S^* mutants

In *Drosophila*, NBs enter a quiescent state during late embryogenesis and remain quiescent until larvae begin feeding (approximately 4 h after larval hatching (Curt *et al*, 2019). Exceptions are the four MB NBs that do not undergo quiescence and continue dividing during late embryo and L1 stages (Ito & Hotta, 1992). To better understand the cause of the observed hyperplasia in *Alk^Y1355S^* larval brains we therefore examined NB quiescence in these mutants. Newly hatched *Alk^Y1355S^* and control (*w^1118^*) L1 larvae brains were dissected and stained for pH3 (Fig. 5A-B’). Similar to controls (*w^1118^*), *Alk^Y1355S^* mutants exhibited four pH3-positive mushroom body NBs, indicating that there are no detectable defects in NB quiescence. Later during early pupal development, NBs shrink, exit the cell cycle and terminally differentiate (Homem *et al*, 2014), and in adult flies there is no detectable NB proliferation. To exclude the possibility that NBs continue to proliferate at later stages in *Alk^Y1355S^* mutants, we measured NB size in pre-pupae of control (*w^1118^*) and *Alk^Y1355S^* in comparison to wandering L3 larvae. Similar to control (*w^1118^*), *Alk^Y1355S^* NBs undergo a progressive size reduction in pre-pupal stages indicating proper terminal differentiation (Suppl. Fig. 6A-B). Finally, we did not detect excessive proliferation in adult brains of *Alk^Y1355S^* mutants using anti-PH3 staining (Suppl. Fig. 6C). Taken together, no perturbations in NB quiescence were observed in *Alk^Y1355S^* mutants in L1, excluding this mechanism as the source of the observed hyperplasia. In addition, no change in proliferation dynamics could be detected at adult stages.

**Figure 5.**
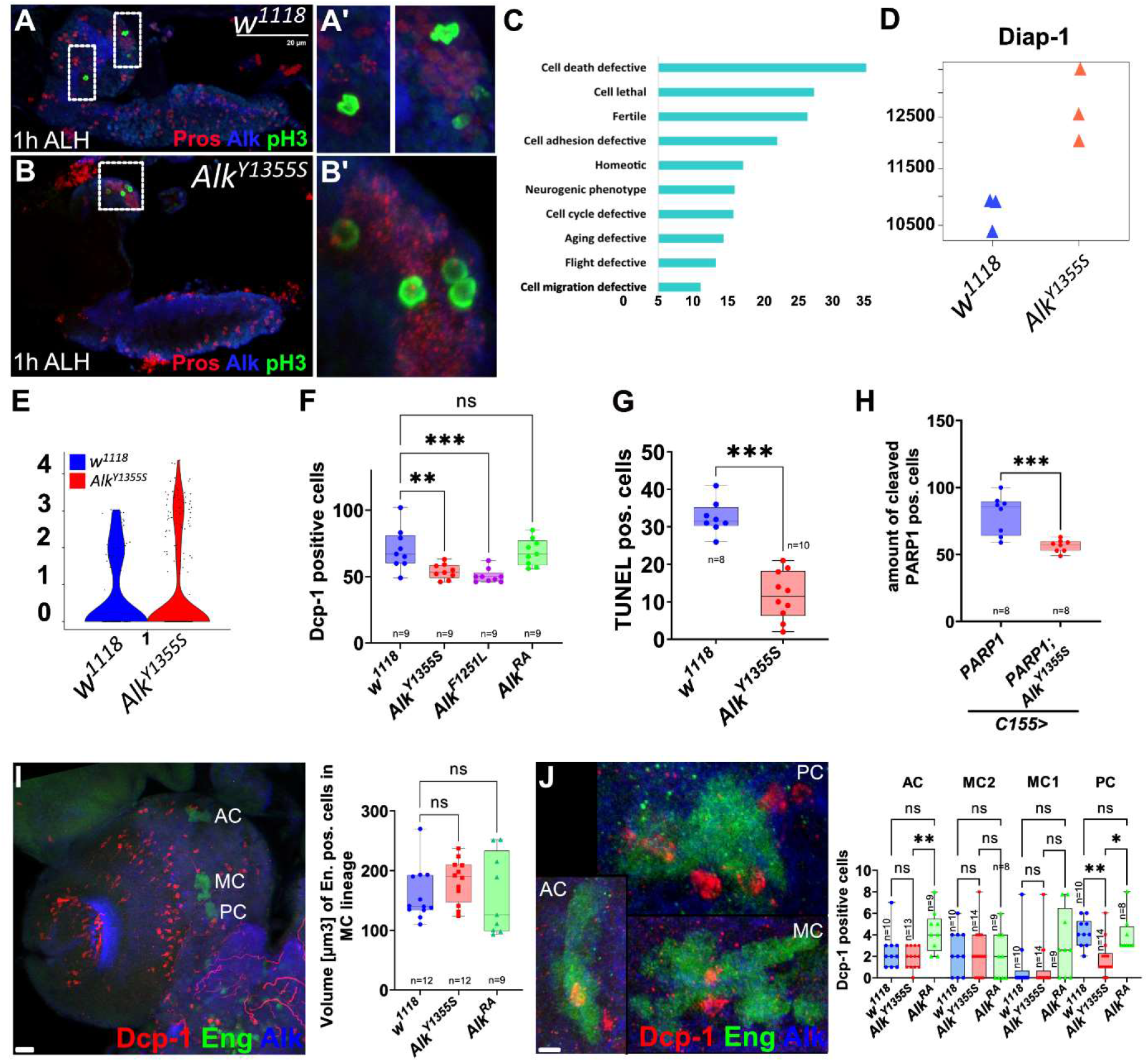
*Alk^Y1355S^* larval brains exhibit decreased levels of apoptosis. **A-B** Four pH3-positive MB NBs are detected in both *Alk^Y1355S^* (A) and *w^1118^* control (B) first instar larval brains. Scale bar 20 µm. **C** Gene Ontology (GO) analysis of genes enriched in sub-cluster 1 of the reanalyzed neuroblast enriched cells cluster in *Alk^Y1355S^*, highlighting an enrichment in cell death defective components in the *Alk^Y1355S^* scRNA-seq dataset. **D** Expression of *Diap1* in bulk RNASeq from both control and *Alk^Y1355S^* larval brains. **E** Violin plot indicating *Diap1* expression in the neuroblast enriched cells population in the *Alk^Y1355S^* and *w^1118^* controls. **F** *Drosophila* Cleaved caspase-1 staining (Dcp-1) shows a significant decrease in Dcp-1-positive cells in the central brain area of *Alk^Y1355S^*. One-way Anova, Dunnett’s multiple comparison test, p>0.001. **G** TUNEL analysis indicates decreased numbers of apoptotic cells in *Alk^Y1355S^* compared to *w^1118^* controls. Unpaired t test; p<0.001, n=8 (*w^1118^*) and n=10 _(*Alk*_*_Y1355S_*_)._ **H** PARP1 reporter assay in control and *Alk^Y1355S^* reveals a significant decrease in reporter activity in *Alk^Y1355S^* larval brains. Unpaired t test; p<0.001, n=8 for each genotype. **I** Volume analysis of the engrailed-positive MC sub-lineage shows no difference in *Alk^Y1355S^* and *Alk^ΔRA^* compared to the *w^1118^*controls. Scale bar 20 µm. One-way Anova, Kruskal-Wallis test, p=not significant (ns). **J** Analysis of Dcp-1-positive neurons adjacent to the engrailed sub-lineages anterior cluster (AC), posterior cluster (PC) and medial cluster (MC). Graph indicates a significant difference in Dcp-1-positive cells around the AC in *Alk^Y1355S^* and *Alk^ΔRA^,* and around the PC between *Alk^Y1355S^* and w1118 controls. Scale bar 5 µm. One-way Anova, Kruskal-Wallis test. AC p<0.007, MC1 p=not significant (ns), MC2 p=not significant (ns), PC p<0.001.

### *Alk* mutants display reduced levels of apoptosis

In models of childhood neuroblastoma, expression of the MYCN oncogene in the developing neural crest drives tumorigenesis. In a zebrafish neuroblastoma model, the additional transgenic expression of the *Alk^F1174L^* activated variant, overcomes excessive apoptosis observed in MYCN only tumors, decreasing cell death and promoting tumorigenesis (Zhu *et al*, 2012), suggesting an important role for Alk signaling in promotion of survival. Moreover, the zebrafish Alk family RTK Leucocyte tyrosine kinase (Ltk) that functions in neural crest-derived iridiophore development has been reported to promote survival when carrying a neuroblastoma-associated mutation (Fadeev *et al*, 2016). Further, in the fly visual system, Alk has been shown to be important for the survival of L3 neurons (Pecot *et al*., 2014) prompting us to address apoptosis in more detail. Indeed, gene ontology analysis of cluster 1 in our scRNA-seq dataset identified an enrichment of ‘cell death defective’ markers in postembryonic *Alk^Y1355S^* mutant brains (Fig. 5C). These included the Diap1 suppressor of apoptosis, which was also upregulated in bulk RNA-seq of *Alk^Y1355S^* brains (Fig. 5D-E). This prompted us to test whether decreased apoptosis in *Alk^Y1355S^* and *Alk^F1251L^* mutant brains may lead to the observed increased numbers of neurons. To investigate this, Cleaved Caspase (Dcp-1) antibody staining of *Alk^Y1355S^*, *Alk^F1251L^* and *Alk^ΔRA^* L3 larvae brains was performed and Dcp-1 positive cells in the central brain area were counted. Significantly reduced Dcp-1 levels were observed in both *Alk^Y1355S^* and *Alk^F1251L^* mutants, although we did not observe increased apoptosis in *Alk^ΔRA^* mutant larval brains (Fig. 5F). The Dcp-1 analysis was complemented with two additional apoptosis assays, firstly a TUNEL assay and secondly with a PARP1 reporter, which both confirmed reduced levels of apoptosis in the *Alk^Y1355S^* mutants (Fig. 5G-H). Finally, we analyzed apoptosis in the controlled setting of the engrailed-positive AC, PC, MC1, and MC2 hemi-lineages, in which immediate progeny of specific GMCs are removed via apoptosis (Kumar *et al*, 2009). In the MC2 lineage, all engrailed-positive neurons are removed via apoptosis, while engrailed-positive or negative neurons in the MC1 lineage do not undergo programmed cell death (Kumar *et al*., 2009). In the AC and PC lineage the engrailed-negative GMC progeny are removed via apoptosis (Kumar *et al*., 2009). We therefore aimed to see differences in expansion of the MC lineage when Alk signaling is active and potentially suppresses apoptosis in that area. Analysis of the overall volume of the MC lineage via engrailed staining, while simultaneously staining for Dcp-1, failed to identify any differences in MC lineage size in either *Alk^Y1355S^* or *Alk^ΔRA^* compared with control (*w^1118^*) (Fig. 5I). However, analysis of Dcp-1-positive cells within and in close proximity to the other hemi lineages identified significantly more Dcp-1-positive cells around the AC lineage in *Alk^ΔRA^* compared to control (*w^1118^*) and *Alk^Y1355S^.* This finding was supported by the presence of significantly less Dcp-1-positive neurons around the PC lineage in *Alk^Y1355S^* compared to control (*w^1118^*) and *Alk^ΔRA^* (Fig. 5J). Taken together, these four independent assays support decreased levels of apoptosis in *Alk^Y1355S^* mutants as a contributory factor to the increased numbers of surviving neurons observed.

### Perturbed neuronal fate in the mushroom body lineage in *Alk^Y1355S^* mutant brains

Interestingly, scRNA-seq analysis highlighted increased proportions of cells that exhibited high levels of expression of neuronal lineage markers such as *mamo*, *br*, *tlk, nk, DnaJ-1*, *Hr4* and *Hsp68* in cluster 1 of *Alk^Y1355S^* larval brains (Fig. 6A-B). To visualize differentially expressed markers we analyzed the expression of the BTB/POZ-containing, C2H2 zinc finger transcription factor Mamo via antibody staining in control (*w^1118^*) and *Alk^Y1355S^*. Mamo has been reported to be critical for temporal specification of α’β’ mushroom body (MB) neurons (Liu *et al*., 2019; Rossi & Desplan, 2020). Initial investigation confirmed that Mamo is expressed in Alk-positive larval neurons in the MB lineage (Fig. 6C). As our scRNA-seq datasets detected significantly more *mamo* expression in the *Alk^Y1355S^* mutant allele (Fig. 6A-B), we measured the overall volume of Mamo-positive cell clusters within the MB lineage, observing an increase in the total volume of Mamo-positive cells in *Alk^Y1355S^* (Fig. 6D) compared to the control (*w^1118^*), validating our scRNA-seq data.

**Figure 6.**
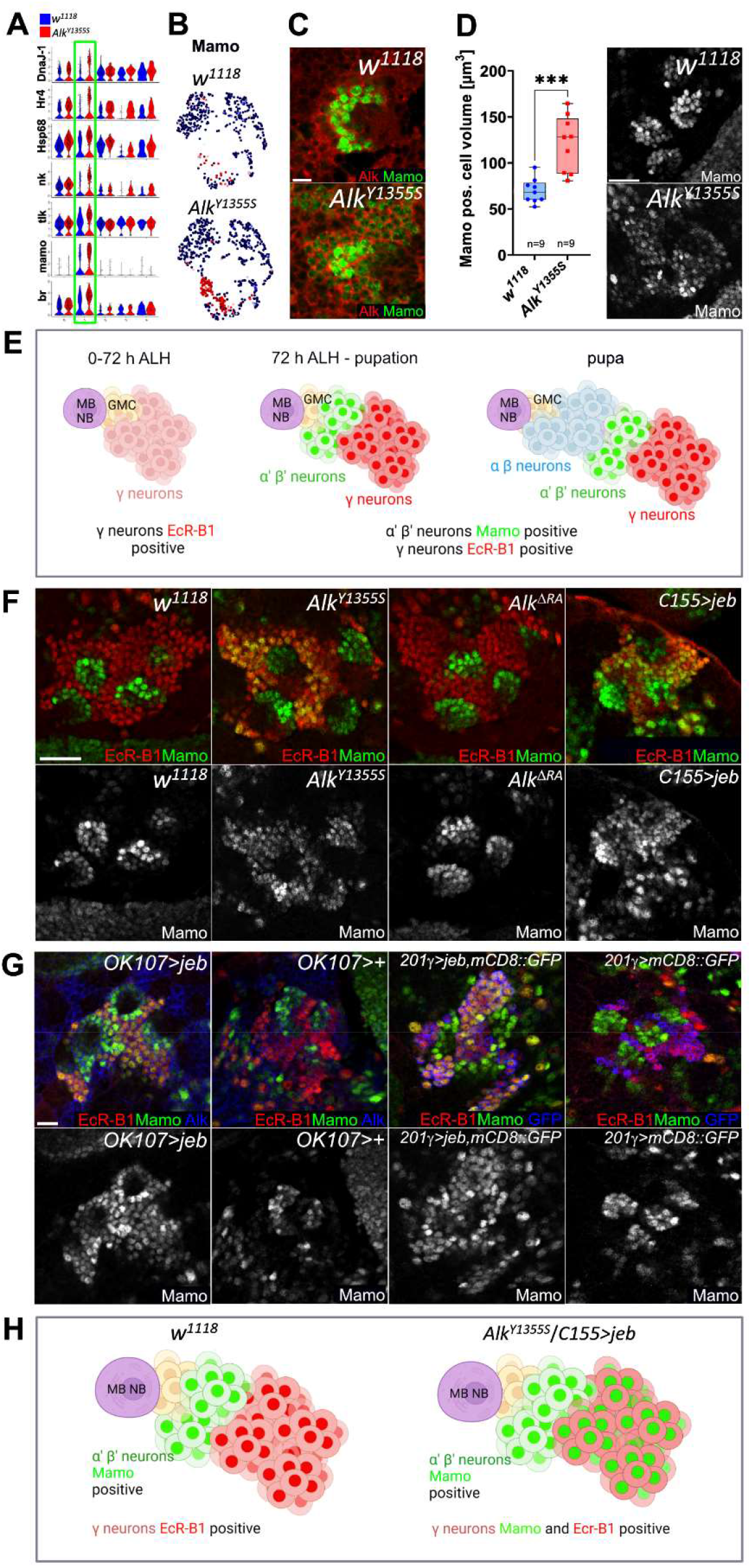
Ectopic Mamo expression in the MB γ lineage in *Alk^Y1355S^* indicates perturbed α’β’ and γ neuronal differentiation. **A** Violin plots indicating expression of *mamo*, *br*, *tlk, nk, DnaJ-1*, *Hr4* and *Hsp68* in cluster 1 of the *Alk^Y1355S^* scRNA-seq dataset. **B** Feature plots projecting *Mamo* mRNA expression in both wild-type and *Alk^Y1355S^* scRNA-seq datasets. **C** Mamo (green) and Alk (red) are co-expressed in neurons. Scale bar 5 µm. **D** Quantification of Mamo-positive cell area in *Alk^Y1355S^* compared to *w^1118^* controls. Unpaired t test, p<0.001. **E** Schematic showing the generation of the three different major MB lineages during development (adapted from (Lee *et al*., 1999)). Schematic created with BioRender.com. **F** Mamo is detected in the MB γ neuron lineage (EcR-B1) in *Alk^Y1355S^*, but not in *w^1118^* controls or the *Alk^ΔRA^* mutant. Overexpression of Jeb ligand (*C155>jeb)* phenocopies the ectopic Mamo expression in the γ lineage observed in *Alk^Y1355S^*. Scale bar 20 µm. **G** Mushroom body-specific *OK107-* and *201γ-Gal4* drivers overexpressing Jeb phenocopy the *Alk^Y1355S^* phenotype. Scale bar 20 µm. **H** Schematic summarizing the ectopic Mamo expression observed in *Alk^Y1355S^*/*C155>jeb* compared to control (*w^1118^*) MB lineages. Schematic created with BioRender.com.

To examine Mamo expression in *Alk^Y1355S^* larval brains further, we focused on the MB NB lineage in wL3 larvae. The MB lineage comprises three major cell types: early born γ neurons, middle born α’β’ neurons and late born αβ neurons (Fig. 6E)(Lee *et al*., 1999). Mamo expression is restricted to the α’β’ neurons whereas EcR-B1 is exclusively expressed within the γ neurons in wL3 (Fig. 6E) (Lee *et al*., 2000; Liu *et al*., 2019; Rossi & Desplan, 2020). Remarkably, we observed ectopic expression of Mamo in Ecdysone receptor (EcR-B1)-positive γ neurons in *Alk^Y1355S^* mutant wL3 brains (Fig. 6F, H). To confirm that excessive Alk activity results in improper Mamo expression we ectopically overexpressed Jeb with *C155-Gal4*, which also resulted in ectopic expression of Mamo in EcR-B1-positive MB γ neurons (Fig. 6F, H). Analysis of Mamo in *Alk^ΔRA^* mutants larval brains showed that although increased Alk activation leads to Mamo expression, Alk is not critically required for normal Mamo expression in the α’ and β’ lineage, as Mamo expression in *Alk^ΔRA^* mutants α’β’ MB neurons is unchanged (Fig. 6F). These findings were confirmed by expression of Jeb using the MB specific *OK107-Gal4 and 201γ-Gal4* drivers, which also led to Mamo expression in wL3 EcR-B1-positive MB γ neurons (Fig. 6G, H).

We next asked whether ectopic Mamo expression leads to a neuronal fate change in γ neurons in *Alk^Y1355S^* larval brains which is maintained in adults, comparing axonal morphology and the MB cell bodies (MBC) in wild type and *Alk^Y1355S^* with antibodies to the Rho guanine exchange factor (RhoGEF) Trio. Trio is important for directional extension of neurites in the MB and is strongly expressed in the cytoplasm and plasma membrane of α’β’ cell bodies, while in γ cell bodies expression is enriched at plasma membranes but is largely absent from the cytoplasm (Awasaki *et al*, 2000). We also examined the Abrupt zinc finger, BTB domain containing transcription factor that functions in motoneuron guidance and connectivity (Hu *et al*, 1995).

In adult flies the γ, α’β’ and αβ MBC branch anteriorly though the peduncle into three main lobes: the bifurcating α’β’and αβ lobes and a single γ lobe (Fig. 7A) (Crittenden *et al*., 1998). No gross defects in overall MB morphology were observed in *Alk^Y1355S^* adult brains (Fig. 7B), however examination of Trio and Abrupt expression in the MBCs identified more cell bodies with enhanced and cytoplasmic Trio expression (Fig. 7C) whereas expression of Abrupt was decreased in *Alk^Y1355S^*. These results were strengthened by increased cytoplasmic Trio positive cell bodies in *OK107>jeb* adults (Fig. 7D), further indicating that increased Alk signaling shifts γ identity to a more α’β’ neuronal identity in larval stages, which is maintained through metamorphosis into adulthood.

**Figure 7.**
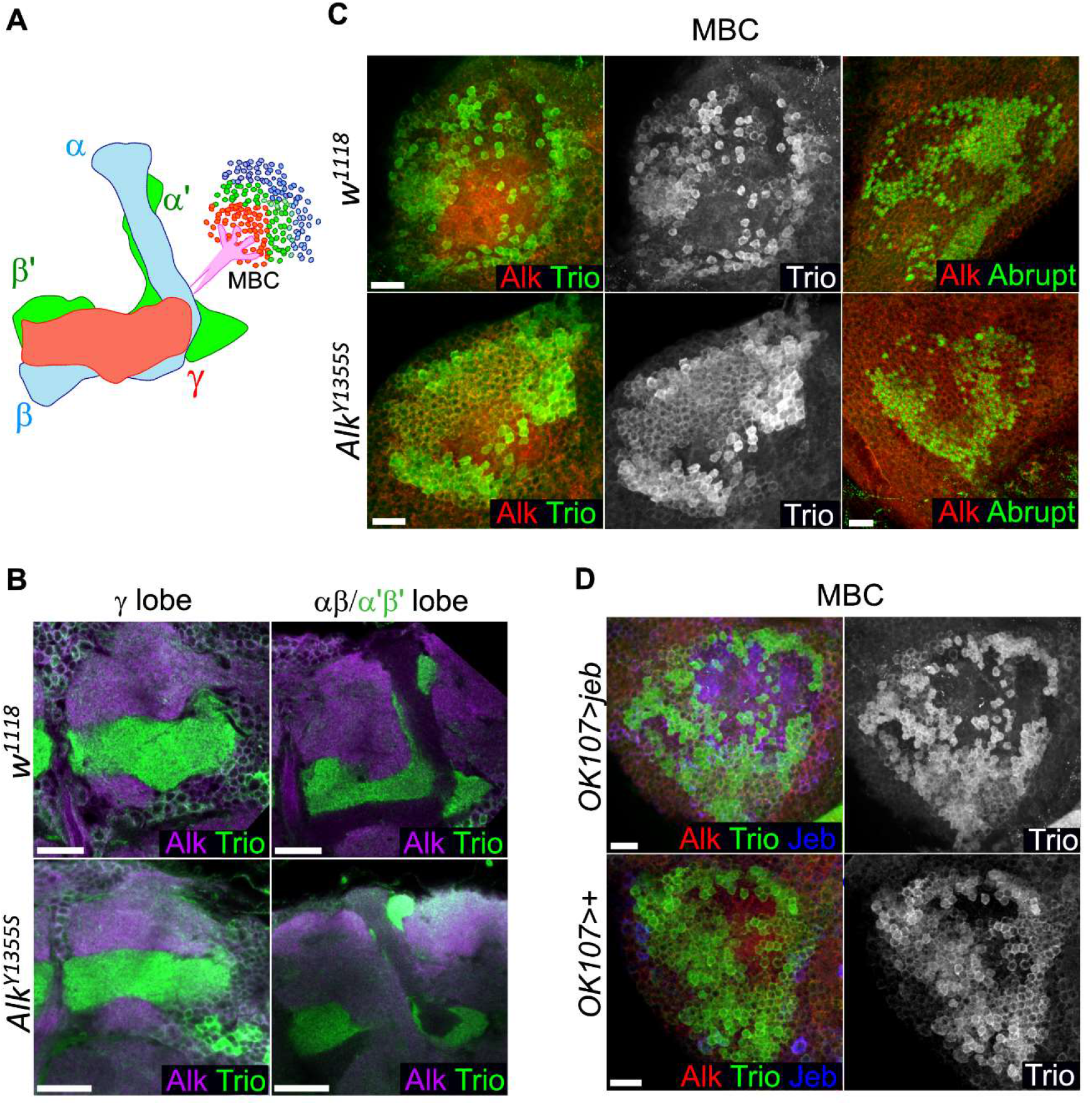
Perturbed neuronal differentiation in *Alk^Y1355S^* is maintained to adulthood. **A** Schematic displaying the three main MB neuronal types (αβ, α’β’, and γ) and the mushroom body cell bodies (MBC) in adults. **B** Analysis of α’β’, and γ lobes using Trio and Alk staining. No gross differences in axonal phenotype could be detected in *Alk^Y1355S^* compared to control (*w^1118^*). Scale bar 20 µm. **C** Analysis of Trio and Abrupt expression in the MBC indicates an increase in Trio-positive MBCs, together with a reduction in Abrupt-positive cell bodies in *Alk^Y1355S^*. Scale bar 20 µm. **D** Trio expression in *OK107>jeb* shows increased Trio-positive cell bodies compared to *OK107>/+* controls. Scale bar 10 µm.

Taken together, our data show that neuroblastoma associated, Alk point mutations do not result in changes in NB number, or in gross increases in proliferation, but rather perturb neuronal fate specification, which leads to precocious Mamo expression in the γ cell lineage of *Alk^Y1355S^* mutant wL3 brains that persist until adult stages.

## Discussion

This work set out to address mechanisms by which point mutations in the Alk receptor tyrosine kinase affect neural development. This is particularly relevant in pediatric neuroblastoma, where patients with mutations in human *ALK* often have poor prognosis, and where tumors are considered to arise due to defects in differentiation during development of the nervous system. For this reason, we generated and characterized two independent *Alk* mutations (*Alk^Y1355S^* and *Alk^F1251L^*) in *Drosophila,* for which the human orthologues (*ALK^Y1278^* and *ALK^F1174L^)* have previously been characterized as “gain-of-function” mutations in neuroblastoma patients (Chand *et al*., 2013; Guan *et al*., 2017).

As in other model systems where Alk mutation has been investigated, including mouse and zebrafish, flies harboring *in locus Alk* modifications are homozygous viable and fertile (Borenas *et al*., 2021b; Cazes *et al*., 2014; Fadeev *et al*., 2016). In keeping with earlier findings where *Drosophila* Alk has been activated by overexpression of the Jeb ligand, *Alk^Y1355S^* and *Alk^F1251L^* mutants are small in size (Gouzi *et al*., 2011; Wolfstetter *et al*., 2017), confirming that these mutations result in increased signaling output through the Alk pathway. Employing *Drosophila* as genetic model allowed us to investigate the process of neurogenesis in a highly controlled environment, revealing that: (i) *Alk^Y1355S^* and *Alk^F1251L^* mutant larval brains exhibit neuronal hyperplasia; (ii) no increase in neuroblast numbers; and (iii) no dramatic increase in neuronal proliferation, suggesting that another mechanism underlies the increase in neurons observed in these mutant alleles. The observation of increased numbers of neurons in either *Alk^Y1355S^* or *Alk^F1251L^* mutant larval brains was also seen when we overexpressed the *Drosophila* Alk ligand Jeb in the CNS and furthermore are in keeping with reports from several labs of hyperplasia in peripheral nervous system ganglia of *Alk* “gain-of-function” mice (Borenas *et al*., 2021b; Cazes *et al*., 2014; Ono *et al*., 2019). Interestingly, these studies suggest that, while the mechanisms underlying this neuronal hyperplasia are not well understood, it is clear that these mutations in the endogenous *Alk* locus collaborate with the MYCN oncogene to drive highly penetrant and aggressive neuroblastoma in mice (Borenas *et al*., 2021b; Cazes *et al*., 2014; Ono *et al*., 2019).

Defects in apoptosis during development can also lead to hyperplasia in the nervous system (Zhu *et al*., 2012), and regulation of apoptosis is important to establish the final number of neurons in the CNS (Pinto-Teixeira *et al*., 2016). Indeed, our results support decreased apoptosis as a contributing mechanism underlying the hyperplastic phenotype observed in *Alk^Y1355S^* and *Alk^Y1355S^* brains. Interestingly, an Alk suppressor screen recently suggested that Alk signaling leads to growth advantages and increases the competitive advantage of cells via transforming JNK activity (Wolfstetter *et al*., 2020). These findings are also in keeping with the ability of RTK downstream signaling to drive survival signaling pathways in multiple systems (Bergmann *et al*, 1998; Bergmann *et al*, 2002; Kurada & White, 1998), and are also supported by elegant work in the fly visual system where Alk is required for L3 neuronal survival in the optic lobe (Pecot *et al*., 2014). Moreover, in zebrafish knock-down or inhibition of Alk also induces apoptosis on the fish hindbrain (Yao *et al*., 2013).

One important aspect to consider from our work is the expression of *Alk* itself. Our scRNA-seq datasets identified strong expression of *Alk* mRNA in neuronal lineages particularly in mature neurons, but not in NBs. Indeed, we could also confirm this at the protein levels by Alk immunostaining of larval brains. This Alk expression profile is in line with our observations of normal cessation of NB proliferation in *Alk^Y1355S^* mutant brains and would fit with an anti-apoptotic effect in neurons later during the differentiation process. Whether *Alk^Y1355S^* mutants are protected from apoptosis throughout development and in the adult brain is not known. However, in zebrafish, aberrant signaling of the Alk family RTK, Ltk, which carries a neuroblastoma related mutation (*Ltk^moonstone^*), promotes survival of neural crest derived iridophores and increases their final number (Fadeev *et al*., 2016). This increase in iridophores number is also observed when the zebrafish Alk family ligands, the Alkals, are ectopically expressed, suggesting that inappropriate wild type Ltk activation by its ligands, can also lead to iridophore survival (Fadeev *et al*., 2018). In addition, our findings that *Alk^Y1355S^* and *Alk^Y1355S^* do not drive uncontrolled proliferation in the *Drosophila* brain support earlier reports in mouse and zebrafish models that harbor activating mutations in the *Alk* locus (Borenas *et al*., 2021b; Cazes *et al*., 2014; Fadeev *et al*., 2016; Ono *et al*., 2019), where mutation of *Alk* alone is insufficient to drive spontaneous tumor development.

An important tool generated in this study is the *Alk^ΔRA^* mutant. The *Alk* locus encodes two transcripts, *Alk-RA* and *Alk-RB*, which differ in their 5’ non-coding regions, employing alternative promoters that encode for the same protein (Loren *et al*., 2001; Mendoza-Garcia *et al*., 2017). From earlier work we know *Alk^ΔRB^* mutant embryos, in which the promoter of the *Alk-RB* isoform has been disrupted, fail to express detectable Alk protein in the embryonic VM and exhibit an *Alk* loss of function phenotype (Mendoza-Garcia *et al*., 2017). We also noted that expression of Alk in the embryonic CNS is not compromised in the *Alk^ΔRB^* mutant. In this work, we show that the *Alk-RA* transcript is expressed in the CNS, and by removing the promoter of this isoform, we have generated an *Alk^ΔRA^* mutant that is viable, but which expresses very low levels of Alk mRNA and protein in the larval CNS. Analysis of this *Alk^ΔRA^* mutant suggests that Alk activity is not critical for neuronal differentiation, and is in agreement with a role for Alk in stress related and sensory responses, such as the brain-sparing response and olfactory functions previously described in the fly (Cheng *et al*., 2011; Gouzi *et al*., 2011). Our observation of increased apoptosis in this *Alk^ΔRA^* mutant, while slight, supports the finding of decreased apoptosis in *Alk^Y1355S^* and *Alk^Y1355S^* brains. Whether higher levels of apoptosis might be observed in conditions of stress or injury in the *Alk^ΔRA^* mutant will be interesting to study in the future.

Our results also produced several surprising findings. One of these was that neither *Alk^Y1355S^* nor *Alk^F1251L^* was able to rescue founder cell specification in the developing VM in the absence of Jeb ligand, suggesting that these activating *Alk* alleles are still ligand dependent. The issue of ligand dependency has not been tested in either mouse, zebrafish or human systems, where ALK is activated by the ALKAL ligands (Fadeev *et al*., 2018; Guan *et al*., 2015; Mo *et al*, 2017b; Reshetnyak *et al*, 2015). Although it is clear that both *Alk^Y1355S^* and *Alk^F1251L^* are ligand-dependent in the developing fly VM, we do not have the tools to test this in the *Drosophila* brain and therefore cannot be sure if the observed ligand dependence is tissue specific. Nonetheless, this finding has interesting and important implications for neuroblastoma. Although human *ALK-F1174L* and *ALK-Y1278S* (orthologous to *Drosophila Alk^F1251L^* and *Alk^Y1355S^*) are classified as ligand independent mutations (Chand *et al*., 2013), we cannot exclude that these human ALK mutations still require ligands to initiate activation at endogenous levels of expression. Thus, ALK ligands might still be important for tumor formation and progression, even in the case of activating ALK mutations. Understanding this will require further investigation in the fly as well as in other model systems. Indeed, this hypothesis could be tested by investigating the effect of ALKAL ligand deletion on ALK/MYCN-driven neuroblastoma in mouse models.

The use of scRNA-seq analysis, in addition to bulk RNA-seq, was ultimately critical in allowing us to identify the subtle effects of *Alk* mutation in the larval brain. To facilitate access to our scRNA-seq data, we have employed the ShinyCell R for interactive interfaces package (Ouyang *et al*, 2021), available at : https://ruthpalmer.shinyapps.io/third__instar_larval_brain/. Importantly, in addition to our finding of defective apoptosis that was identified as enriched in the scRNA-seq dataset, we have also been able to identify defects in neuronal differentiation during development in *Alk^Y1355S^* mutant *Drosophila*. By comparison of scRNA-seq datasets from *Alk^Y1355S^* mutant brains with control, we were able to identify several mis-regulated factors, including *mamo*, which could be experimentally interrogated. The development of the MB, which is comprised of approximately 2000 neurons of three main neuronal types (γ, α’β’ and αβ), in *Drosophila* offered a unique opportunity for this validation (Cognigni *et al*., 2018; Ito *et al*., 1997; Lee *et al*., 1999). These three neuronal types, γ, α’β’ and αβ, are produced sequentially, during defined developmental periods in response to Imp and Syp gradients (Liu *et al*., 2015; Ren *et al*, 2017; Syed *et al*., 2017a; Yang *et al*, 2017). Elegant work has shown that the terminal identity and maintenance of MB α’β’ neurons is established by Mamo expression (Liu *et al*., 2019), and that Mamo expression depends on extrinsic receptor signaling via the activin receptor Babo, as *babo* mutant clones exhibit a loss of α’β’ neurons (Rossini *et al*, 2020). While we were unable to identify MB neuronal differentiation defects in *Alk^ΔRA^* mutants, the γ-neuron lineage was Mamo–positive in *Alk^Y1355S^* mutant wL3 brains, a result that was confirmed by overexpression of the Jeb ligand in the CNS. Thus, aberrant Alk activity leads to inappropriate expression of Mamo in the γ linage at this stage, and perturbation of MB neuronal fate, however, Alk is not required for expression Mamo in wild type α’β’ neurons. We were also able to show that these defects in larval MB neuron development are not transient, but persist to adult stages, where the MB neuron cell bodies of *Alk^Y1355S^* mutants exhibit expanded Trio expression. In mice, Trio plays important roles in the regulation of actin dynamics which is crucial for axonal guidance (Paskus *et al*, 2020; Zong *et al*, 2015). Adult mice lacking Trio exhibit hippocampal abnormalities and decreased brain size together with defects in learning ability (Zong *et al*., 2015). The perturbations in MB lineages we observe in this work are in keeping with the role of the MB in learning and memory, where Alk has previously been reported to have a role in the ability of flies to learn (Gouzi *et al*., 2011).

The perturbation of neuronal differentiation described here as a result of activation of Alk has implications for higher organisms, as well as for human neuroblastoma. Neuroblastoma is thought to develop from an aberrantly developed neural crest derived cell somewhere in vicinity of the adrenal medulla and/or the sympathetic ganglia (Huber *et al*, 2009; Saito *et al*, 2012). During development these cells undergo migration to destinations close to the dorsal aorta where they form the sympathetic ganglia and the chromaffin cells that contribute to the adrenal medulla. Recent work has shed light on the role of the peripheral stem cells known as Schwann cell precursors (SCPs) during mouse embryonic development (Furlan *et al*, 2017). Although we do not know the effect of Alk activation on differentiation in the mouse nervous system, our findings here suggest that the cell fate change/differentiation process may be perturbed, a subject that can be addressed in future work.

Together, the results presented here identify neuronal hyperplasia, together with decreased levels of apoptosis, in the brains of *Alk^Y1355S^* and *Alk^F1251L^* mutant flies. Further, with the aid of scRNA-seq, we could identify molecular components in the neuronal differentiation process that arise as a consequence of *Alk* mutation. Employing the well-characterised process of MB neuron development, we have been able to experimentally validate these neuronal differentiation defects, and additionally show that they persist to adult stages. To our knowledge, this is the first time that molecular defects in neuronal differentiation have been identified in response to *Alk* activating mutations. These findings have potentially important implications for pediatric neuroblastoma, where mutation of human ALK at orthologous sites (*ALK-Y1278S* and *ALK-F1174L*), is associated with the progression of these tumors in the developing peripheral nervous system.

## Material and Methods

### Drosophila husbandry

Standard *Drosophila* husbandry procedures were followed (Ashburner, 1989). *Drosophila* stocks were reared on standard diet at room temperature. Crosses were performed at 25°C, 60% humidity and 12/12 hours day-night cycle.

### Fly stocks

Fly stocks obtained from the BDSC (NIH P40OD018537): *P{GawB}elav^C155^ (C155-Gal4, BL-458)*, *y^1^, {Mvas-Cas9}ZH-2A, w^1118^* (BL-51323), *w^1118^; P{70FLP}10* (BL-6938), *w^1118^; P{GAL4-Act5C(FRT.CD2).P}S, P{UAS-RFP.W}3/TM3, Sb^1^*(Bl-30558), *w*; P{Ubi-p63E(FRT.STOP)Stinger}15F2*(BL-32251), y1 *\*; P{UAS-CD8::PARP1-Venus}3* (BL-65609), *w*; P{GawB}OK107 ey^OK107^/In(4)ci^D^, ci^D^ pan^ciD^ sv^spa-pol^* (BL-854*), y^1^ w^67c23^; P{UAS-mCD8::GFP.L}LL5 P{GawB}Tab2^201Y^* (BL-64296). Other lines used were as follows: *C155-Gal4*, *UAS-Alk.EC (Bazigou et al., 2007)*, *UAS-Jeb* (Varshney & Palmer, 2006), *jeb^weli^* (Stute et al., 2004), *UAS-hALK.F1174L*, *UAS-hALK* (Martinsson *et al*, 2011), *UAS-hALK.Y1278S* (Guan *et al*., 2017), UAS-GFP.caax (Finley *et al*, 1998).

### Generation of *Alk^F1251L^*, *Alk^Y1355S^* and *Alk^ΔRA^* employing CRISPR/Cas9-mediated genome editing

The ***Alk^F1251L^*** and ***Alk^Y1355S^*** alleles were generated using *CRISPR/Cas9* (Gratz *et al*, 2013). sgRNA target sites were identified using the flyCRSIPR target finder (https://flycrispr.org/)(Gratz *et al*, 2014). Two sgRNA for each approach (sgRNA1 Y1355S: TCGCCGATTTTGGCATGTCCCGG, sgRNA2 Y1355S: TCGGACTACTATCGCAAGGGAGG; sgRNA1 F1251L: CCTGAAGG AGGCGGCCATAA TGG, sgRNA2 F1251L: CAAG<u>T</u>TCA ATCACCCGAATA TGG) were cloned into pBFv-U6.2 expression vector (Genome Engineering Production Group at Harvard Medical School). sgRNAs and donor constructs were injected into *y^1^, {Mvas-Cas9}ZH-2A, w^1118^* embryos by BestGene Inc.. Mutations were inserted by homology directed repair (HDR) using a donor construct (Gratz *et al*., 2014). HDR donor constructs were cloned into pBluescript II by Genescript. Design of the *Alk^Y1355S^* and *Alk^F1251L^* donor constructs was as follows: *Y1355S*: containing silent mutations to prevent annealing and destruction of the donor by Cas9: TtGCtGAcTTcGGtATGTCaCG and TCaGAtTAtTAcCGtAAaGGtGG. Homology arms: 507 bp upstream; 506 bp downstream of the desired mutation site (Suppl. Methods Fig. 1). *F1251L*: silent mutations plus integrated nucleotide exchange to create desired mutation: CTaAAaGAaGCaGCaATcATG GCaAAG **cTt** AAcCAtCCaAAcATG, homology arms: 1052 bp upstream, 902 bp downstream (Suppl. Methods Fig. 2). Positive *CRISPR/Cas9* mutants were identified by PCR screening of single males using primers directed against the region of HDR integrated silent mutations (forward primer ATC CTA ATG ATC TCG CTT GCC GTG, Y1355S reverse primer: 5’-CTcCCGATCaGAtTAtTAcCGtAAaG-3’ and F1251L reverse primer: 5’-TT tGGaTGgTTaAgcTTtGC CAT gAT-3’) and subsequent sequencing (GATC Biotech).

To generate the kinase dead ***Alk^D1345A^*** mutant allele and the ***Alk^D1345A, Y1355S^*** double mutant, the *Alk^Y1355S^* HDR donor construct was modified accordingly using the Q5® site-directed mutagenesis kit (NEB, #E0554S). Due to the close proximity of both codons within the sequence, the guide RNA target sequences (sgRNA Y1355S nr.1 and nr.2, cloned into *pU6-BbsI-chiRNA*)(Gratz *et al*., 2013) could be used for injection into *y^1^, {Mvas-Cas9}ZH-2A, w^1118^* embryos. Positive candidates were obtained by single-fly PCR-screening with Y1355S forward and reverse primers (*Alk^D1345A, Y1355S^*) or lethality screening (*Alk^D1345A^*). All positive candidates obtained failed to complement the *Alk^KO^* kinase domain deletion mutation.

The ***Alk^ΔRA^*** mutant allele was created with CRISPR/Cas9 exploiting the NHEJ repair pathway by a dual guide approach (sgRNA10_RA 5’- CCGTCTATCCGCGATTCTGAGGG-3’; sgRNA20_RA 5’- GCGACAGTGGCGCACTCTGGCGG-3’). This mutant was designed to specifically generate a deletion within the 5’UTR of *Alk-RA* isoform. sgRNA sequences were cloned into the pBFv-U6.2 expression vector (Genome Engineering Production Group at Harvard Medical School), and the subsequent constructs injected into *vasa (vas)-Cas9* (BL-51323) embryos by BestGene Inc.. Screening of deletion events was performed on F1 single males by PCR using primers flanking the intended deletion (forward primer: *Alk-RA Fw1* 5’-cgaaatttttcctgcagctc-3’; reverse primer: *Alk-RA Rv1* 5’-atggggtccttaatgcactg-3’) and deletions were further characterized by sequencing (GATC Biotech).

### Pupal size measurement

6 virgin females and 4 males of each genotype were reared for two days on standard diet at 25°C. The vials were kept at 25°C till pupae collection. Shortly before hatching female pupae were collected on double sided tape on a slide, frozen at −20°C and then analyzed with a ZEISS AxioZoom V16 stereo microscope.

### Fixation and immunohistochemistry staining of embryos

Embryo staining was carried out after (Muller, 2008). For phospho-ERK staining we followed the protocol from (Gabay *et al*, 1997).

### Fixation and immunohistochemistry staining of larval and adult brains

Brains of all stages were dissected in PBS and collected in PBS containing 3.7 % formaldehyde on ice and subsequently fixed in the same solution for 30 min at RT followed by permeabilization in PBS containing 1% Triton-x-100 for 10 min and three washes in PBS containing 0.5% triton-x-100 (PBT). Samples were blocked in PBT; 5 % goat serum (Jackson Immuno Research) in PBT, then primary antibody added overnight at 4°C. Samples were washed 4 times in PBT and incubated in PBT containing the secondary antibody and DAPI for 2 hours at RT followed by 4 washes in PBT. Samples were mounted in Fluoromount G and analyzed on a ZEISS Axio Imager.72 microscope. A ZEISS LSM800 confocal microscope and ZEN Blue edition software were used to acquire images.

Primary antibodies: gp-anti-Alk (1:1000), rb-anti-Alk (1:1000)(Loren *et al*., 2001), gp-anti-Jeb (1:1500)(Englund *et al*., 2003), m-anti-activated MAPK/dephosphorylated ERK1/2 (1:200, Sigma-Aldrich #M8159), rb-anti-pH3 (1:500, Millipore, #0657C) rb-anti-Miranda (1:1000, Abcam #197788), ch-anti-GFP (1:500, Abcam #ab13970), rb-anti-RFP (1:1000, Abcam #ab62341), rb-anti-cleaved PARP1 (1:50, Abcam #ab2317, anti-β-Galactosidase (1:150, Abcam #ab616), rb-anti-Dcp-1 (1:50; Cell Signaling Technology #9578), Developmental Studies Hybridoma Bank: m-anti-Dlg (1:500; #4F3), rat-anti-ElaV (1:100; #9F8A9), m-anti-EcR-B1 (1:100; #ADA4.4), m-anti-Trio (1:100; # 9.4A), m-anti-Abrupt (1:50). Antibodies gifted: guinea pig-anti-Mamo (1:100; from Claude Desplan lab (Rossi & Desplan, 2020)), rb-anti-Ase (1:400; from Cheng-Yu Lee lab (Weng *et al*, 2010)). Secondary antibodies were purchased from Jackson Immuno Research. DAPI (1mg/ml) (Sigma-Aldrich, #D9564-10MG) was used at 1:1000.

### In situ hybridization

In situ hybridization was carried out as described previously (Mendoza-Garcia *et al*., 2017).

### Hybridization chain reaction (HCR) v.3

The HCR probe sets were generated by Molecular Instruments (www.molecularinstruments.com) against full length *Alk* and *jeb* mRNA sequences from the NCBI database (Accession number *jeb*: NM_136882; accession nr Alk: NM_001274098.1). HCR amplifier for *jeb*: B3 with the A488 amplifier fluorophore, *Alk*: B2 with the 546 amplifier fluorophore. Fluorescent *in situ* hybridization was carried out according to the protocol for *Drosophila* embryos for HCR v.3 from Molecular Instruments with the following modifications: Proteinase K digestion was carried out with 0.01 mg/ml proteinase K in PBST for 10 min.

### Immunoblotting

Brains from 30 wandering third instar larvae were dissected, collected on ice and subsequently lysed in RIPA buffer. Protein concentration was measured using the BCA-kit (Thermo Scientific). Laemmli buffer (final concentration 1x) was added prior to loading on a 7.5% SDS/PAGE gel. After transfer to PVDF (Millipore, #IPVH00010), membranes were immunoblotted with primary antibodies overnight at 4°C. Primary antibodies used: gp-anti-Alk (1:3000) (Loren *et al*., 2001) and rb anti-tubulin (1:5000; Cell Signaling, #11H10). After washing, membranes were incubated with goat anti-rb HRP (Invitrogen, #11859140) secondary antibodies at RT for 1 hour. ECL prime (Thermo Scientific, # GERPN2236) was used for detection.

### Clonal analysis

To generate RFP or GFP clones, *hs-flp* females were crossed to males carrying *Act5c-Gal4>FRT.CD2>UAS-RFP* (Pignoni & Zipursky, 1997) or *Ubi-p68E>FRT.stop>stinger* in either control (*w^1118^*) *or Alk^Y1355S^* backgrounds. L1 Larvae were collected from 2-hour collection plates after hatching and grown at 25°C for 72 hours. Larvae were subsequently heat shocked at 37°C for 10 min and reared at 25°C for a further 24 hours.

### EdU pulse and chase experiments

The Click-iT™ EdU Alexa Fluor™ 488 Imaging Kit from Thermofisher(Catalog) was employed according to manufacturer’s instructions. L3 larvae were reared on 0.2M EdU in potato-based fly food for 3 hours (3h pulse) for analysis of NB I, 1.5 hours (1.5h pulse) for NB II and dissected immediately afterwards, or reared for 24 hours at 25°C (24h chase, for NB I) and then dissected and fixed. Staining was carried out according to the manufacturer’s instructions.

### TUNEL assay

ApopTag® Fluorescein In Situ Apoptosis Detection Kit (Sigma-Aldrich, catalog number S7110) was employed according to the manufacturer’s instructions.

### *In vitro* brain culture

Wandering L3 larvae were washed twice in PBS and once in 70% ethanol before collection and dissection in Schneideŕs medium. Dissected brains were cultured in Schneideŕs Medium containing 10% heat activated FBS (Sigma, # F7524), 0.01% Insulin solution (Sigma, I0516-5ML), 1% Penicillin/Streptavidin (HyClone, # SV30010), 1 µg/ml 20-hydroxy-ecdysone (Sigma, H5142-5MG) for 18, 36 or 120 hours. Prior to fixation brains were washed 3 times in PBS and then fixed in 4% formaldehyde in PBS for 20 min RT. Staining was caried out as described above in the immunohistochemistry staining section.

### Brain dissociation for cell sorting and scRNA-seq analysis

Wandering third instar larvae were washed in PBS, and brains dissected and collected in ice-cold PBS for a maximum of 1 h. After rinsing twice in ice-cold PBS, brains were incubated in 200 µl of dissociation solution (1 mg/ml collagenase I) for 1 h with continuous agitation, after which the reaction was stopped by addition of 10 ml of PBS (0.04% BSA) and left for 10 min to precipitate larger debris. The resulting dissociated cell solution was sieved through 70 µm and 40 µm cell strainers to remove cell clumps. The filtered suspension was centrifuged at 3000 rpm for 5 min at 4 °C. Supernatant was discarded and cells resuspended at 10^8^ cells/ml (or at least in 100 μl) of PBS 2% FBS and 1 mM CaCl_2_. Dead cells or debris from the dissociated samples were removed using the EasySep Dead Cell Removal (Anexin V) Kit (STEMCELL, #17899) according to the manufacturer’s guidelines. The remaining cells were respectively labelled with aqua-fluorescent reactive dye (dying cells) and calcein violet AM (living cells) using the LIVE/DEAD Violet Viability/Vitality Kit (Molecular Probes, #L34958) under manufacturer’s guidelines. Finally, each sample was washed twice in PBS 2% fetal bovine serum and resuspended in 500 µl PBS 2% fetal bovine serum. Living cells were enriched using a FACSAria III cell sorter (BD biosciences) based on LIVE/DEAD staining. Cells were sorted using an 85 µm nozzle into Eppendorf tubes that had been pre-coated with PBS containing 2% BSA.

### Single cell library preparation, sequencing and scRNA-seq aAnalysis

Approximately 7000 sorted cells were directly loaded in sheath fluid onto one lane of a Chromium 10X chip (10X Genomics) and libraries prepared using the normal workflow for Single Cell 3’ v3 libraries (10X Genomics) and sequenced on the NextSeq 500 platform (Illumina). Initial preprocessing of scRNA-seq datasets began with demultiplexing, read QC, mapping (STAR aligner with the *Drosophila melanogaster* genome) and quantification of barcodes and UMIs using the Cell Ranger (5.0) software. The wild-type dataset resulted in a total of 4,081 cells and the *Alk^Y1355S^* resulted in a total of 4,222 cells. Downstream analysis including cell preprocessing, count-normalization, feature selection, integration, clustering, dimensionality reduction/projection, trajectory inference and differential expression testing was performed using the R-based pipeline Seurat (4.0.3) (Stuart *et al*, 2019) and the python-based pipeline Scanpy (Wolf *et al*, 2018). Cell quality was assessed by the proportion of cells with unique feature counts between 200 and 5000 (removing poor quality cells), and transcripts between 500 and 50,000 (removing low-quality transcripts) and with less than 20% mitochondrial genes. After preprocessing 3967 and 4099 cells remained with a total of 10,411 and 10,316 RNA features, for the wild type and *Alk^Y1355S^* datasets, respectively. Normalization of scRNA-seq datasets and integration analysis were performed with the SCTransform approach (Hafemeister & Satija, 2019). The top 3000 highly variable genes were selected to determine the true dimensionality of the dataset. PCA was used for clustering and based on Elbow-Plots, the number of PCs were determined. Non-linear dimensional reduction using UMAP (Uniform Manifold Approximation and Projection) and neighborhood identification with k=30 was used. Clusters were identified using the Louvain algorithm (Blondel *et al*, 2008), with 0.56 resolution. To determine clusters, marker genes of each cluster were identified by FindAllMarkers function (Seurat) and logistic regression analysis (Scanpy). Based on canonical markers and gene expression profiles, genes similar clusters were merged and annotated based on existing knowledge and literature (Ariss *et al*, 2020; Brunet Avalos *et al*, 2019; Cattenoz *et al*, 2016; Estacio-Gomez *et al*, 2020; Michki *et al*, 2021).

To identify differentially expressed genes, expression profiles of each cluster across *Alk^Y1355S^* and wild type conditions were compared by the MAST approach (Finak *et al*, 2015), with a threshold of log fold-change of average expression between the two groups >= 0.2 and P_value <= 0.05. Cluster correlation was performed using Pearson correlation algorithms (Scanpy), and relationship between clusters determined by unrooted-phylogenetic tree (Seurat). Gene Ontology (GO) analysis was performed using the R package, EnrichR (employing FlyEnrichr as reference) (Kuleshov *et al*, 2016), on Cluster 1 marker genes identified from Neuroblast Enriched Cells (*Alk^Y1355S^*).

### RNA-sequencing and analysis

3rd instar larvae (40 - 45) were collected and quickly washed in water to remove food and yeast. Larvae were placed in ice-cold PBS in a depression well. Fine forceps were used to dissect larval CNS’ into Eppendorf tubes containing 250 μl of ice-cold solution (equal volume of both RNA free PBS and RNALater solution from Invitrogen). Afterwards, the tubes containing CNS were centrifuged at 2000 rpm for 5 min at 4 °C and supernatant removed. 500 μl fresh RNALater solution was added, and samples were stored at −80°C. Once CNS collection for 3 biological replicates per genotype was obtained, RNA-extraction was carried out according to the manufacturer’s protocol (Promega ReliaPrepTM RNA Tissue Miniprep System, #Z6111). RNA concentration was measured using NanoDrop OneC (Thermo Scientific) and RNA-integrity was checked by gel electrophoresis. 4-6 µg of total RNA/biological replicate sequenced by Novogene Co. Ltd (UK). Samples were assessed for quality with the Agilent 2100 Bioanalyzer system and paired-end sequencing performed on an Illumina platform. Over 40 million reads/genotype were generated and mapped to the genome at a rate of over 94-96%. *Drosophila melanogaster* (ensemble bdgp6_gca_000001215_4 genome assembly) was used. HISAT2 algorithm for alignment and DESeq2 R package (Anders & Huber, 2010) for differential gene expression was used.

### Alk domain sequence and structure analysis

Pairwise alignment of Alk domain sequence of *Drosophila Melanogaster* with Human ALK was performed using EMBOSS Needleman-Wunsch algorithm (Madeira *et al*, 2019). The three-dimensional structure of the *Drosophila* Alk kinase domain was modeled based on the human ALK crystal structure dataset template (NCBI Protein Domain and Macromolecular Structures database, PBD ID: 4TT7) and the annotated *Drosophila* Alk-PA sequence (NP_001261027.1) using PyMol 1.8.6.0 software.

### Statistical analysis

Data acquirements and volume calculation was carried out with Microsoft Office Excel software. Statistical analysis was performed with GraphPad Prism 9 software.

### Data availability

ScRNA-seq and bulk RNA-seq datasets for this study have been deposited in Gene Expression Omnibus (GEO) with accession numbers GSE198850 and GSE198812, respectively.

## Acknowledgments

We thank Cheng-Yu Lee and Claude Desplan for sharing fly stocks and reagents. We acknowledge Bloomington Drosophila Stock Center (NIH P40OD018537) for fly stocks used in this study. The ElaV, Trio, Abrupt, Dlg, Eng, EcR-B1 antibodies developed by G. Rubin, C. Hama, S. Crews, C. Goodman (Dlg/Eng) and C. Thummel respectively, were obtained from the Developmental Studies Hybridoma Bank, created by the NICHD of the NIH and maintained at The University of Iowa, Department of Biology, Iowa City, IA 52242. This work has been supported by grants from the Swedish Cancer Society (RHP CAN18/0729; BA CAN200270P), the Children’s Cancer Foundation (RHP 2019-0078; KP TJ2019-0071), the Swedish Research Council (RHP 2019-03914; MB 2019-01708), the Swedish Foundation for Strategic Research (RB13-0204), the Göran Gustafsson Foundation (RHP2016), the Åke Wiberg Foundation (GW M19-0561) and the Knut and Alice Wallenberg Foundation KAW 2015.0144). Author contributions: K.P. and R.H.P. conceived and developed the project. V.A. carried out the bioinformatics analysis with input from K.P.. K.P., G.W., P.M.G., T.M., B.A. and M.B. performed the wet lab experiments. R.H.P. supervised the project. K.P. and R.H.P. wrote the first manuscript draft with input from V.A. and G.W.. All authors contributed to the final version of the manuscript. We thank Bengt Hallberg, Ezgi Uçkun and Sanjay Kumar Sukumar for critical reading of the manuscript.

## Competing interests

The authors declare that they have no competing interests.

## Supplementary Figure Legends

**Figure S1.**
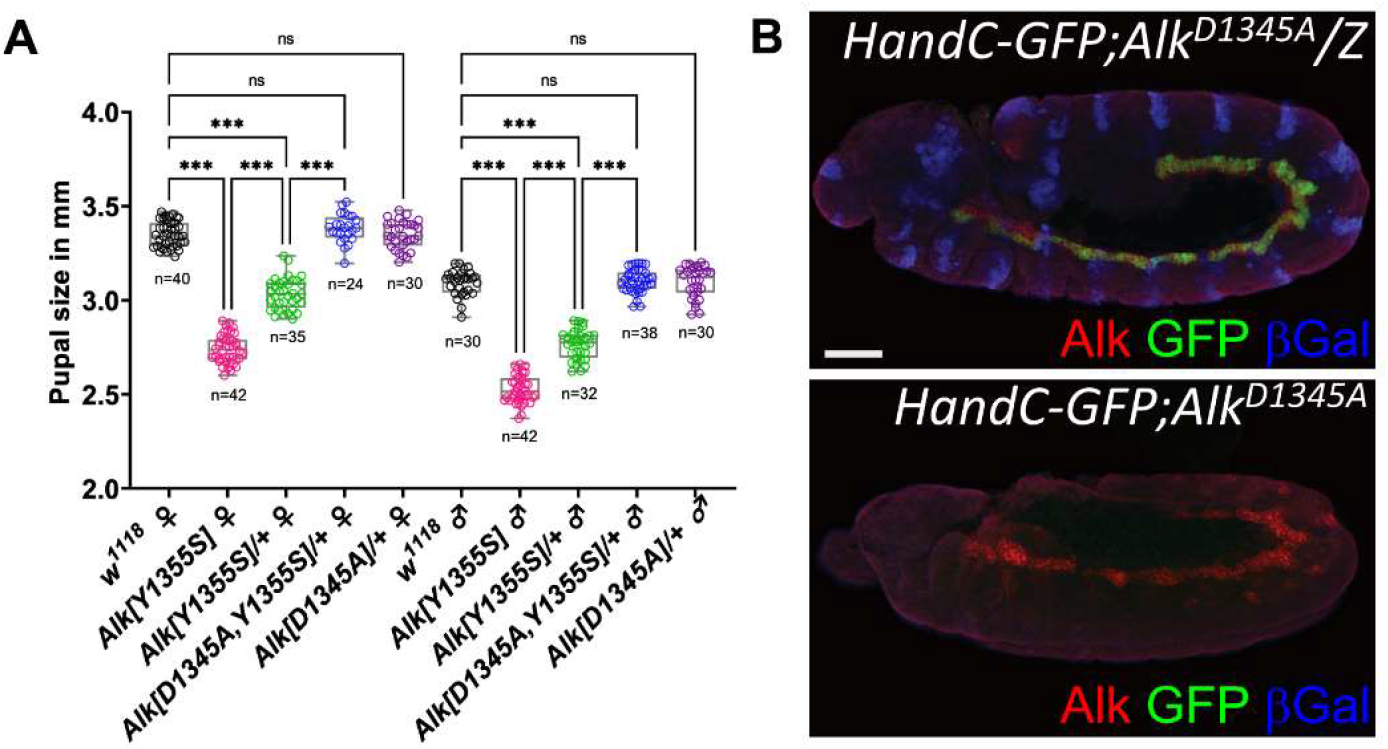
Analysis of *Alk^D1345A^* and *Alk^D1345A, Y1355S^*. **A** Pupal size analysis shows that the small pupal size phenotype of *Alk^Y1355S^/+* is fully rescued in *,Alk^D1345A, Y1355S^/+*,. Ordinary one-way Anova, and Unpaired t test. p<0.001. **B** *Alk^D1345A^* mutants display an *Alk* loss of function phenotype in the VM as visualized by lack of phospho-ERK staining in FCs at stage 10/11. Scale bar 50 µm.

**Figure S2.**
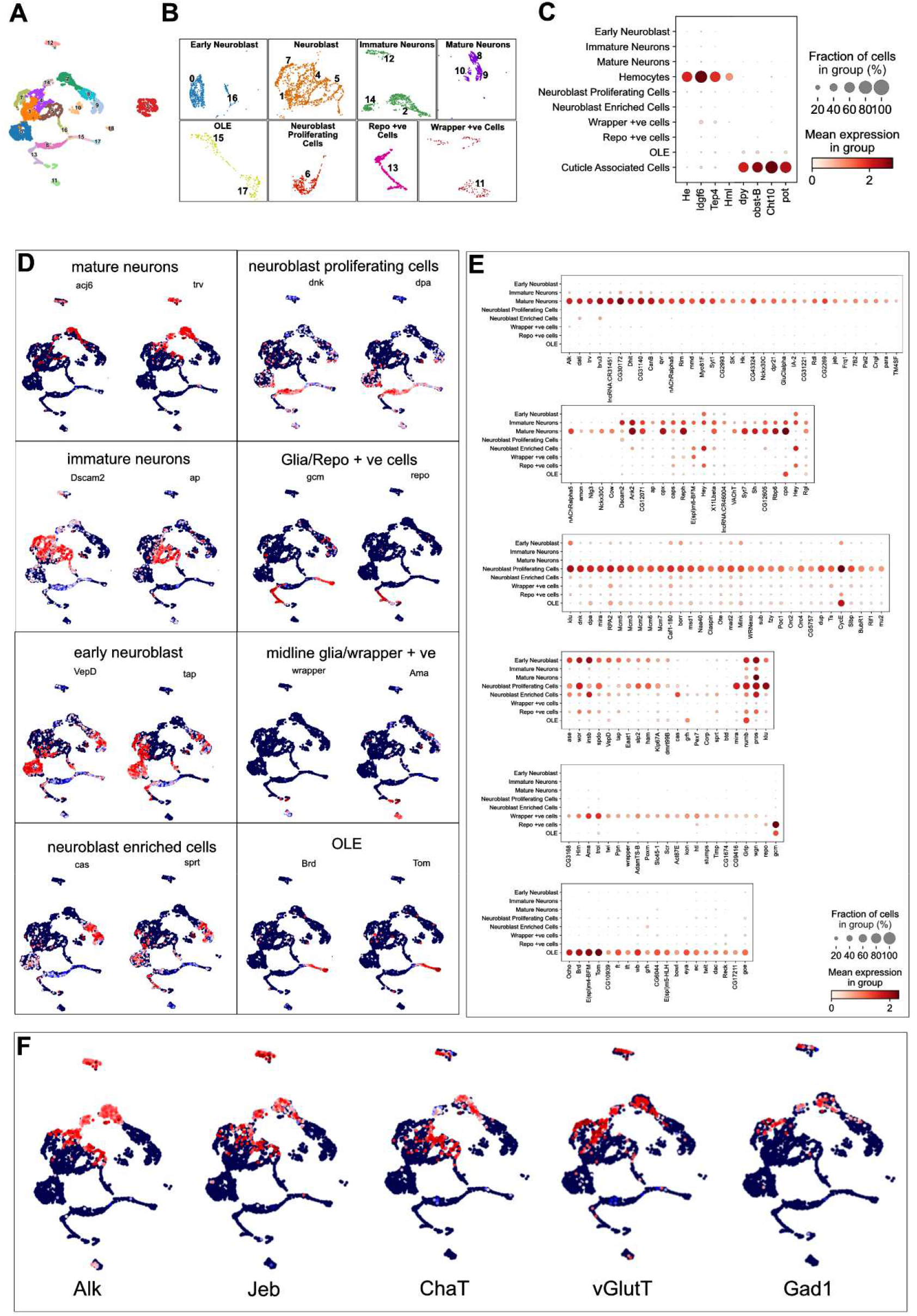
Detailed analysis of scRNA-seq dataset annotation. **A** UMAP displaying the initially defined 19 clusters from the wL3 scRNA-seq dataset. **B** Similar clusters were merged to define the cellular heterogeneity, resulting in eight cell clusters (Early Neuroblast, Neuroblast Enriched Cells, Immature Neurons, Mature Neurons, Neuroblast Proliferating Cells, OLE, Repo +ve cells and Wrapper +ve cells) shown by UMAP. **C** Dotplot showing the expression of canonical markers for the Hemocyte and Cuticle Associated Cell clusters. **D** Feature plots visualizing a pair of canonical markers in the eight clusters. **E** Dotplots indicating unique markers for the eight defined clusters. Color scale indicates mean expression (red gradient) and percentage of cells distributed (dot size). **F** Feature plots showing the expression of *Alk, Jeb*, cholinergic (*ChAT*), glutamatergic (*VGlut*) and GABAergic neurons (*Gad1*) neurons in the wL3 scRNA-seq dataset.

**Figure S3.**
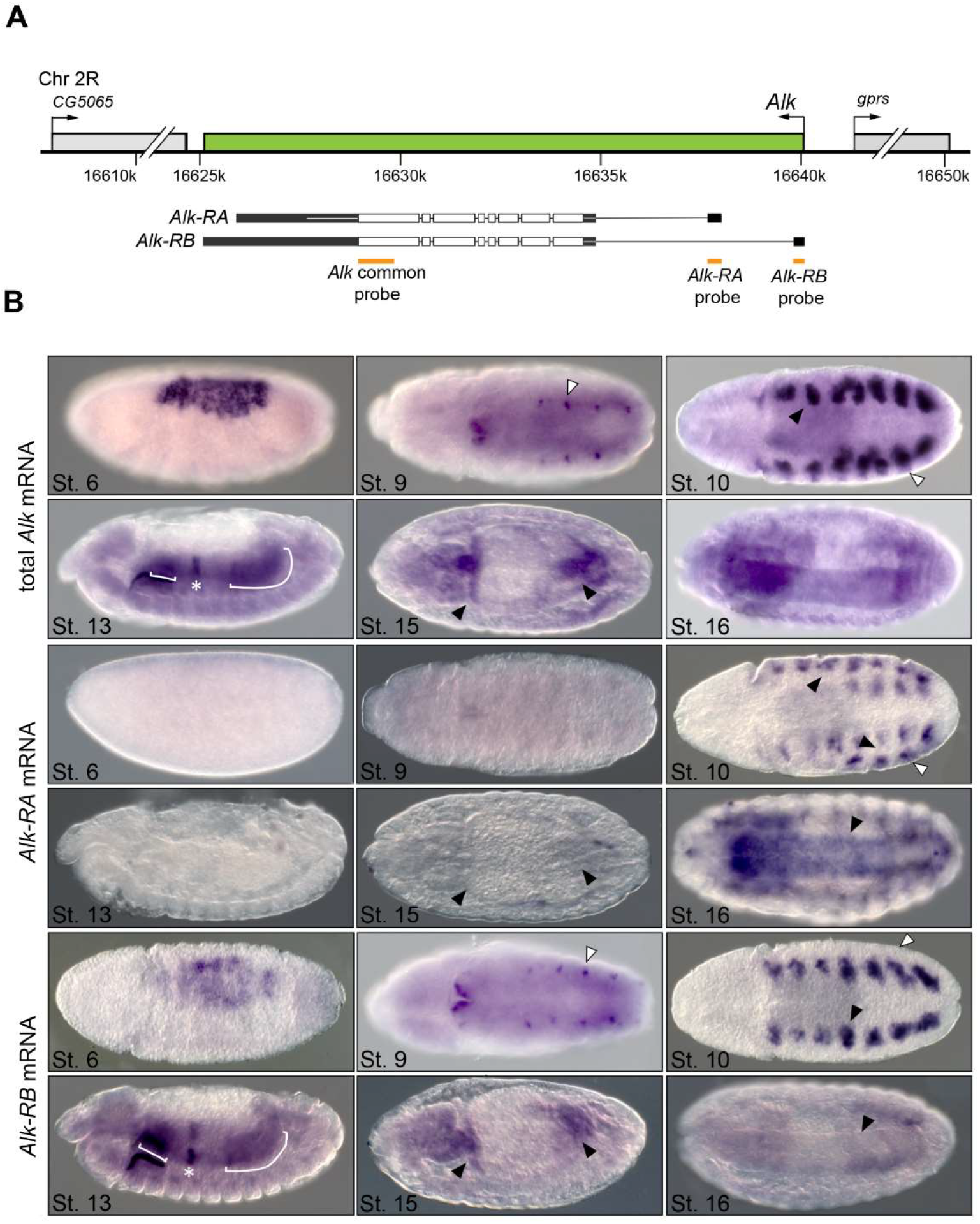
Generation and characterization of the *Alk^ΔRA^* mutant. **A** Schematic outlining the genomic organization of the *Alk* locus (green) and including the neighboring genes *CG5065* and *gprs* (light gray). Intron-exon structure of *both Alk-RA* and *Alk-RB* transcripts is shown below (*Alk* open reading frame in white). Probes employed for *in situ* detection of common *Alk* mRNA, or *Alk-RA/Alk-RB* mRNA transcript specific expressions are indicated in orange. **B** *In situ* showing expression of the common *Alk*, *Alk-RA* and *Alk-RB* transcripts during embryogenesis. Total *Alk* mRNA expression in wild-type embryos. *Alk* transcripts are observed in the amnioserosa (stage 6), the trunk visceral mesoderm (VM) (stage 10, *closed arrowhead*) and the epidermis (stages 10, *open arrowhead*). *Alk* transcripts are present in the visceral muscle, particularly in PS7 (stage 13, *asterisk*) and PS3 (stage 15, *closed arrowhead*). Transcription of *Alk* is also observed in the developing CNS (stage 16). Expression of *Alk-RA* mRNA is observed in the epidermis in close proximity to the trunk VM (stage 10, *open arrowheads*) as well as strongly in the CNS (stage 16). *Alk-RB* mRNA is expressed in the amnioserosa (stage 6), as well as in the trunk VM (stage 10, *closed arrowhead*). The *Alk-RB* transcript is also observed at later stages in the VM, in PS7 (stage 13, *asterisk*) and PS3 (stage 15, *closed arrowhead*). Expression of the *Alk-RB* transcript was not detected in the embryonic CNS (stage 16, *closed arrowhead*).

**Figure S4.**
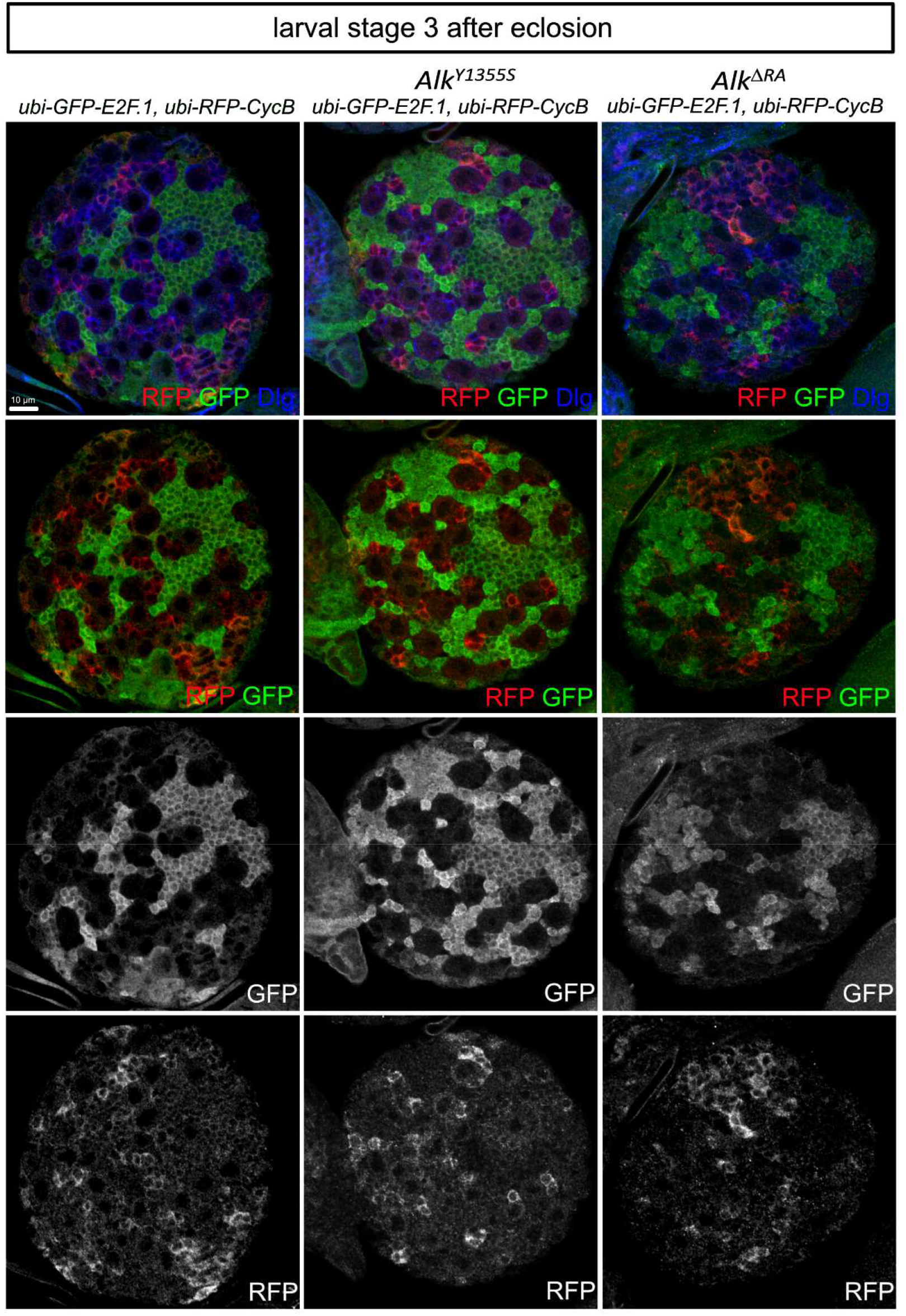
FUCCI system employed in *wild type*, *Alk^Y1355S^* and *Alk^ΔRA^* at L3 stage after eclosion. Fly-FUCCI reporters *ubi-GFP::E2f1* (green) and *ubi-mRFP::CycB* (red) denote G0/1- and S-phase cells, respectively. No gross differences in proliferation can be observed. Dlg2 shown in blue. Scale bar 5 µm.

**Figure S5.**
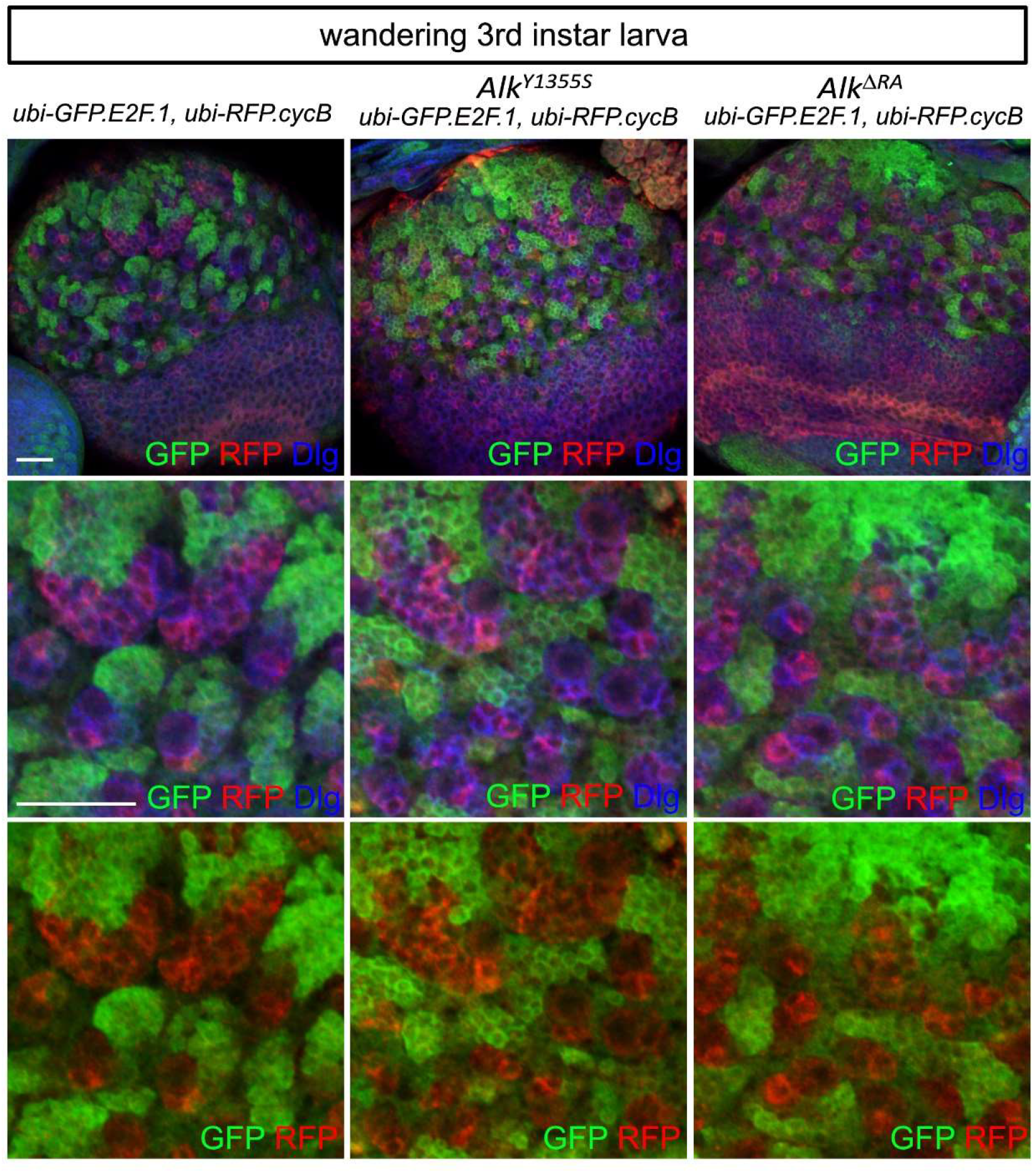
FUCCI system employed in *wild type*, *Alk^Y1355S^* and *Alk^ΔRA^* in wandering 3^rd^ instar larvae. Fly-FUCCI reporters *ubi-GFP::E2f1* (green) and *ubi-mRFP::CycB* (red) denote G0/1- and S-phase cells, respectively. No differences in proliferation in wandering 3^rd^ instar larvae can be observed. Dlg2 shown in blue. Scale bar 20 µm.

**Figure S6.**
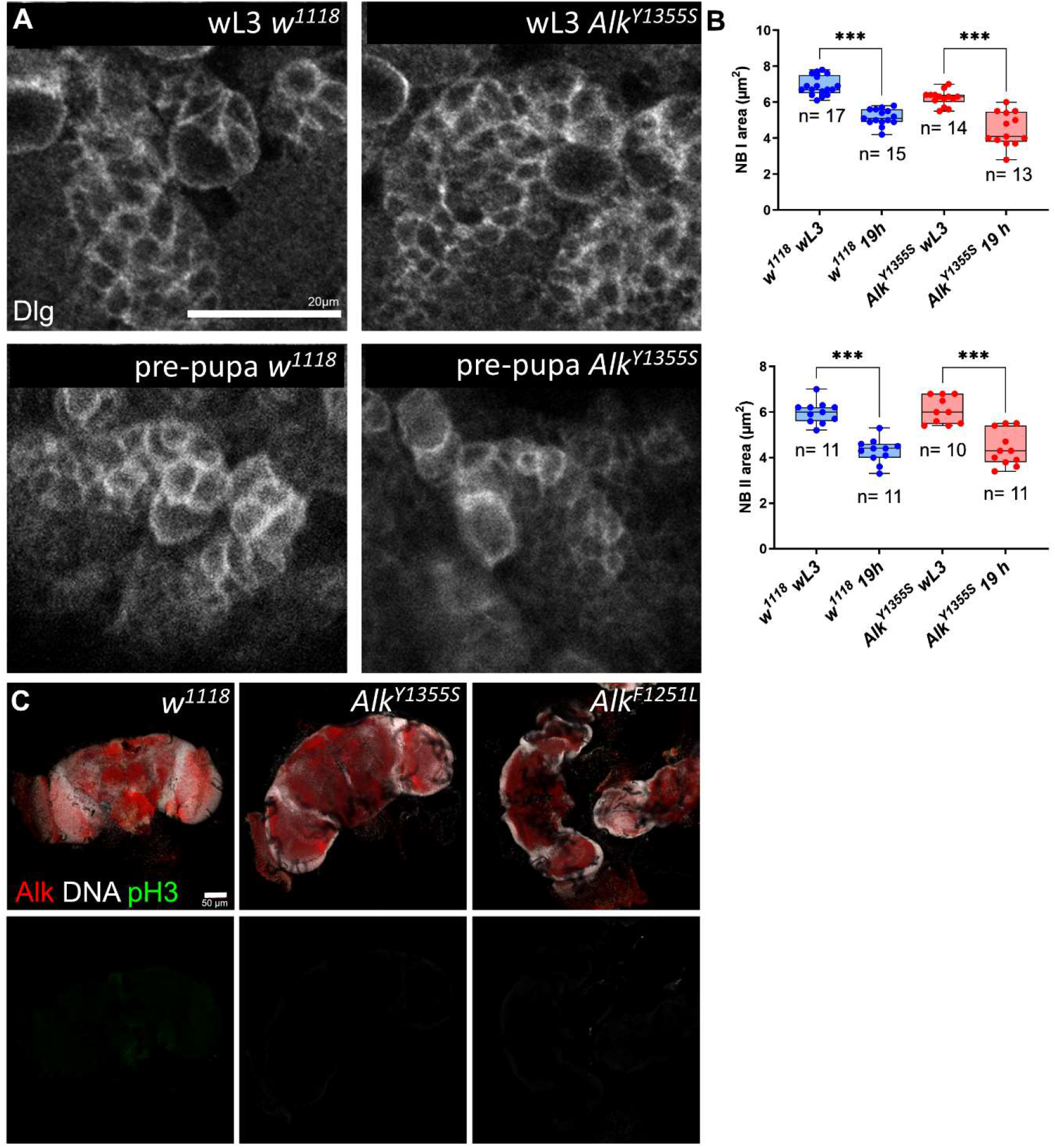
Analysis of proliferation and NB quiescence in wild type and *Alk^Y1355S^* adult brains. **A-B** Type I and II NBs undergo shrinkage from wL3 until 19 h in *in vitro* brain culture. One-way Anova. p<0.001, (n=11; *Alk^Y1355S^* wL3 n=10). Scale bar 20 µm, **C.** No proliferation (pH3) can be detected in adult fly brains in wild type, *Alk^Y1355S^* and *Alk^F1251L^*. Scale bar 50 µm.

## Method supplements

**Method supplement Figure 1.**
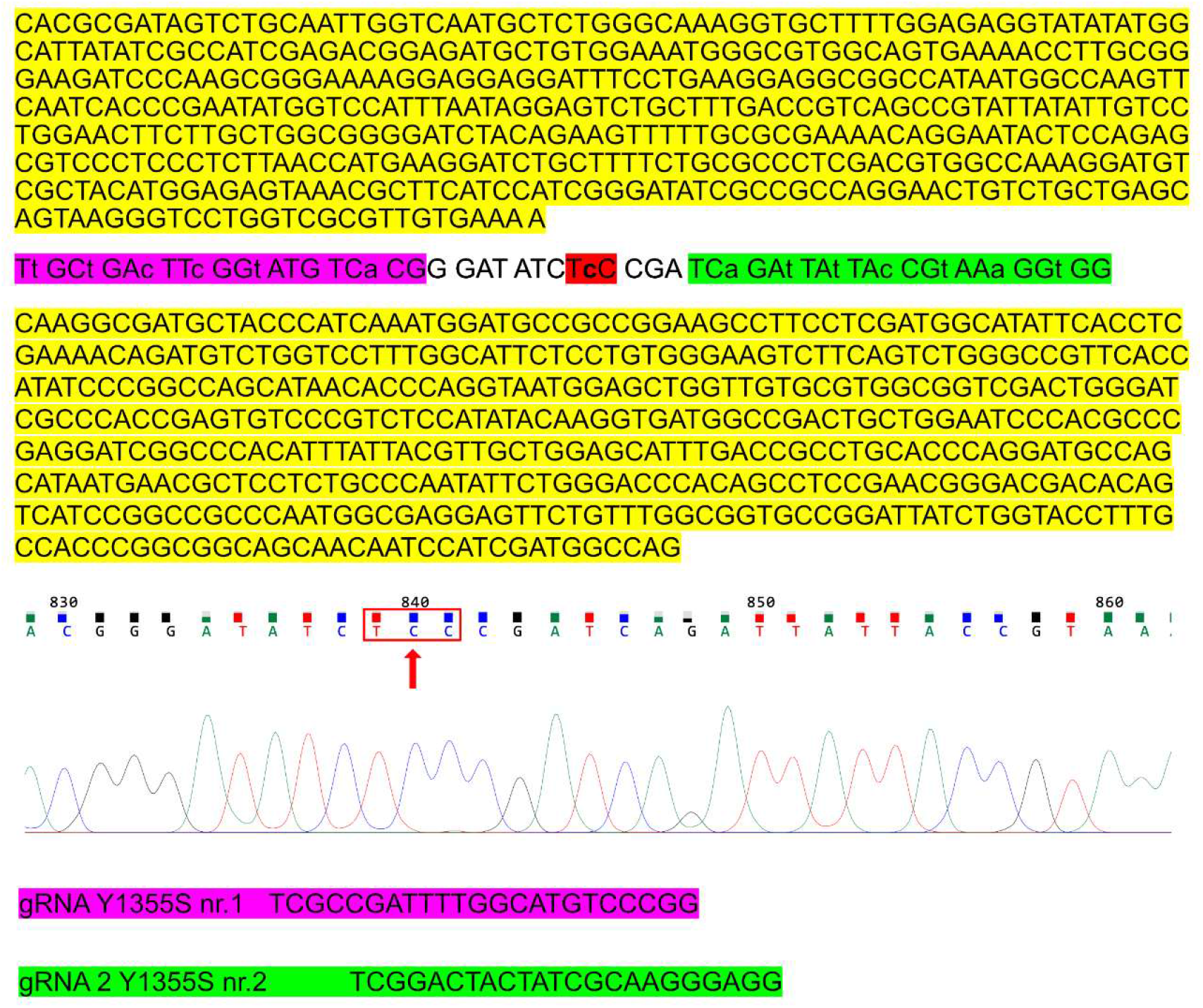
*Alk^Y1355S^* donor construct sequence. Donor construct design: pink and green regions in donor sequence containing silent mutations to prevent the donor construct from destruction during homology induced repair (HDR). Homology arms in yellow. The homology arms are integrated into the endogenous locus during HDR. Chromatogram file from sequenced *Alk^Y1355S^* mutant showing the desired mutation (red arrow), resulting in a change of amino acid from tyrosine to serine (serine, red box). Guide RNA in pink and green.

**Method supplement Figure 2.**
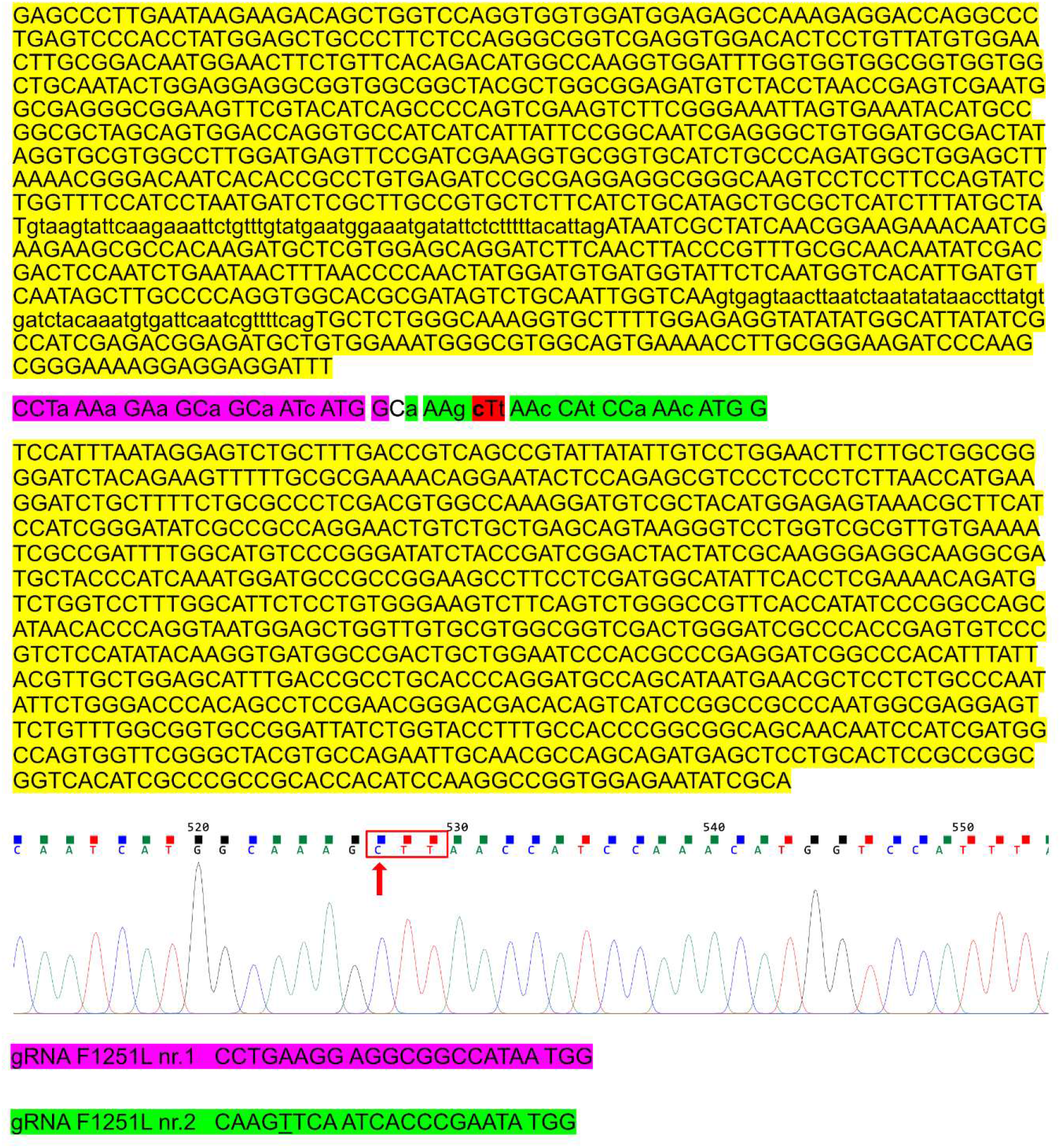
*Alk^F1251L^* mutant donor construct sequence. Donor construct design: pink and green regions containing silent mutations in donor sequence to prevent the donor construct from destruction during homology induced repair (HDR). In red the changed codon. The homology arms are integrated into the endogenous locus during HDR. Chromatogram file from sequenced *Alk^F1251L^* mutant showing the desired mutation (red arrow) resulting in a change of amino acid from phenylalanine to leucine (serine, red box). Guide RNA in pink and green.

**Method supplement Figure 3.**
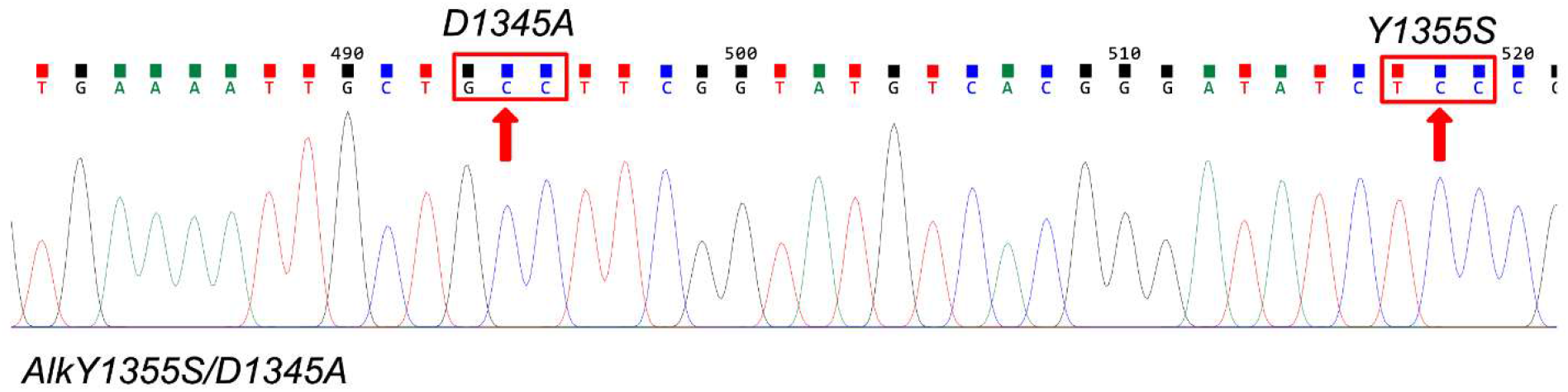
Sequence confirmation of the *Alk^D1345A, Y1355S^* double mutant. Chromatogram showing sequence of the *Alk^D1345A, Y1355S^* double mutant. Red boxes highlight modified codons, red arrows indicate mutated nucleotide.

## Notes

The authors disclose no potential conflicts of interest

## References

Anders S, Huber W (2010) Differential expression analysis for sequence count data. Genome Biol 11: R106

Ariss MM, Terry AR, Islam A, Hay N, Frolov MV (2020) Amalgam regulates the receptor tyrosine kinase pathway through Sprouty in glial cell development in the Drosophila larval brain. J Cell Sci 133

Awasaki T, Saito M, Sone M, Suzuki E, Sakai R, Ito K, Hama C (2000) The Drosophila trio plays an essential role in patterning of axons by regulating their directional extension. Neuron 26: 119–131

Bai L, Sehgal A (2015) Anaplastic Lymphoma Kinase Acts in the Drosophila Mushroom Body to Negatively Regulate Sleep. PLoS Genet 11: e1005611

Bazigou E, Apitz H, Johansson J, Loren CE, Hirst EM, Chen PL, Palmer RH, Salecker I (2007) Anterograde Jelly belly and Alk receptor tyrosine kinase signaling mediates retinal axon targeting in Drosophila. Cell 128: 961–975

Bergmann A, Agapite J, McCall K, Steller H (1998) The Drosophila gene hid is a direct molecular target of Ras-dependent survival signaling. Cell 95: 331–341

Bergmann A, Tugentman M, Shilo BZ, Steller H (2002) Regulation of cell number by MAPK-dependent control of apoptosis: a mechanism for trophic survival signaling. Dev Cell 2: 159–170

Berry T, Luther W, Bhatnagar N, Jamin Y, Poon E, Sanda T, Pei D, Sharma B, Vetharoy WR, Hallsworth A et al (2012) The ALK(F1174L) mutation potentiates the oncogenic activity of MYCN in neuroblastoma. Cancer Cell 22: 117–130

Bilsland JG, Wheeldon A, Mead A, Znamenskiy P, Almond S, Waters KA, Thakur M, Beaumont V, Bonnert TP, Heavens R et al (2008) Behavioral and neurochemical alterations in mice deficient in anaplastic lymphoma kinase suggest therapeutic potential for psychiatric indications. Neuropsychopharmacology 33: 685–700

Blondel VD, Guillaume J-L, Lambiotte R, Lefebvre E (2008) Fast unfolding of communities in large networks. Journal of Statistical Mechanics: Theory and Experiment 2008: P10008

Borenas M, Umapathy G, Lai WY, Lind DE, Witek B, Guan J, Mendoza-Garcia P, Masudi T, Claeys A, Chuang TP et al (2021a) ALK ligand ALKAL2 potentiates MYCN-driven neuroblastoma in the absence of ALK mutation. The EMBO journal 40: e105784

Borenas M, Umapathy G, Lai WY, Lind DE, Witek B, Guan J, Mendoza-Garcia P, Masudi T, Claeys A, Chuang TP et al (2021b) ALK ligand ALKAL2 potentiates MYCN-driven neuroblastoma in the absence of ALK mutation. EMBO J: e105784

Brunet Avalos C, Maier GL, Bruggmann R, Sprecher SG (2019) Single cell transcriptome atlas of the Drosophila larval brain. Elife 8

Caren H, Abel F, Kogner P, Martinsson T (2008) High incidence of DNA mutations and gene amplifications of the ALK gene in advanced sporadic neuroblastoma tumours. Biochem J 416: 153–159

Cattenoz PB, Popkova A, Southall TD, Aiello G, Brand AH, Giangrande A (2016) Functional Conservation of the Glide/Gcm Regulatory Network Controlling Glia, Hemocyte, and Tendon Cell Differentiation in Drosophila. Genetics 202: 191–219

Cattenoz PB, Sakr R, Pavlidaki A, Delaporte C, Riba A, Molina N, Hariharan N, Mukherjee T, Giangrande A (2020) Temporal specificity and heterogeneity of Drosophila immune cells. The EMBO journal 39: e104486

Cazes A, Lopez-Delisle L, Tsarovina K, Pierre-Eugene C, De Preter K, Peuchmaur M, Nicolas A, Provost C, Louis-Brennetot C, Daveau R et al (2014) Activated Alk triggers prolonged neurogenesis and Ret upregulation providing a therapeutic target in ALK-mutated neuroblastoma. Oncotarget 5: 2688–2702

Chand D, Yamazaki Y, Ruuth K, Schonherr C, Martinsson T, Kogner P, Attiyeh EF, Maris J, Morozova O, Marra MA et al (2013) Cell culture and Drosophila model systems define three classes of anaplastic lymphoma kinase mutations in neuroblastoma. Dis Model Mech 6: 373–382

Chen Y, Takita J, Choi YL, Kato M, Ohira M, Sanada M, Wang L, Soda M, Kikuchi A, Igarashi T et al (2008) Oncogenic mutations of ALK kinase in neuroblastoma. Nature 455: 971–974

Cheng LY, Bailey AP, Leevers SJ, Ragan TJ, Driscoll PC, Gould AP (2011) Anaplastic lymphoma kinase spares organ growth during nutrient restriction in Drosophila. Cell 146: 435–447

Cognigni P, Felsenberg J, Waddell S (2018) Do the right thing: neural network mechanisms of memory formation, expression and update in Drosophila. Curr Opin Neurobiol 49: 51–58

Crittenden JR, Skoulakis EM, Han KA, Kalderon D, Davis RL (1998) Tripartite mushroom body architecture revealed by antigenic markers. Learn Mem 5: 38–51

Curt JR, Yaghmaeian Salmani B, Thor S (2019) Anterior CNS expansion driven by brain transcription factors. Elife 8

Doe CQ (2008) Neural stem cells: balancing self-renewal with differentiation. Development 135: 1575–1587

Dong W, Gao YH, Zhang XB, Moussian B, Zhang JZ (2020) Chitinase 10 controls chitin amounts and organization in the wing cuticle of Drosophila. Insect Sci 27: 1198–1207

Englund C, Loren CE, Grabbe C, Varshney GK, Deleuil F, Hallberg B, Palmer RH (2003) Jeb signals through the Alk receptor tyrosine kinase to drive visceral muscle fusion. Nature 425: 512–516

Estacio-Gomez A, Hassan A, Walmsley E, Le LW, Southall TD (2020) Dynamic neurotransmitter specific transcription factor expression profiles during Drosophila development. Biol Open 9

Evans CJ, Liu T, Banerjee U (2014) Drosophila hematopoiesis: Markers and methods for molecular genetic analysis. Methods 68: 242–251

Fadeev A, Krauss J, Singh AP, Nusslein-Volhard C (2016) Zebrafish Leucocyte tyrosine kinase controls iridophore establishment, proliferation and survival. Pigment Cell Melanoma Res 29: 284–296

Fadeev A, Mendoza-Garcia P, Irion U, Guan J, Pfeifer K, Wiessner S, Serluca F, Singh AP, Nusslein-Volhard C, Palmer RH (2018) ALKALs are in vivo ligands for ALK family receptor tyrosine kinases in the neural crest and derived cells. Proc Natl Acad Sci U S A

Finak G, McDavid A, Yajima M, Deng J, Gersuk V, Shalek AK, Slichter CK, Miller HW, McElrath MJ, Prlic M et al (2015) MAST: a flexible statistical framework for assessing transcriptional changes and characterizing heterogeneity in single-cell RNA sequencing data. Genome Biol 16: 278

Finley KD, Edeen PT, Foss M, Gross E, Ghbeish N, Palmer RH, Taylor BJ, McKeown M (1998) Dissatisfaction encodes a tailless-like nuclear receptor expressed in a subset of CNS neurons controlling Drosophila sexual behavior. Neuron 21: 1363–1374

Furlan A, Dyachuk V, Kastriti ME, Calvo-Enrique L, Abdo H, Hadjab S, Chontorotzea T, Akkuratova N, Usoskin D, Kamenev D et al (2017) Multipotent peripheral glial cells generate neuroendocrine cells of the adrenal medulla. Science 357

Gabay L, Seger R, Shilo BZ (1997) MAP kinase in situ activation atlas during Drosophila embryogenesis. Development 124: 3535–3541

George RE, Sanda T, Hanna M, Frohling S, Luther W, 2nd, Zhang J, Ahn Y, Zhou W, London WB, McGrady P et al (2008) Activating mutations in ALK provide a therapeutic target in neuroblastoma. Nature 455: 975–978

Gouzi JY, Bouraimi M, Roussou IG, Moressis A, Skoulakis EMC (2018) The Drosophila Receptor Tyrosine Kinase Alk Constrains Long-Term Memory Formation. J Neurosci 38: 7701–7712

Gouzi JY, Moressis A, Walker JA, Apostolopoulou AA, Palmer RH, Bernards A, Skoulakis EM (2011) The receptor tyrosine kinase Alk controls neurofibromin functions in Drosophila growth and learning. PLoS Genet 7: e1002281

Gratz SJ, Cummings AM, Nguyen JN, Hamm DC, Donohue LK, Harrison MM, Wildonger J, O’Connor-Giles KM (2013) Genome engineering of Drosophila with the CRISPR RNA-guided Cas9 nuclease. Genetics 194: 1029–1035

Gratz SJ, Ukken FP, Rubinstein CD, Thiede G, Donohue LK, Cummings AM, O’Connor-Giles KM (2014) Highly specific and efficient CRISPR/Cas9-catalyzed homology-directed repair in Drosophila. Genetics 196: 961–971

Guan J, Umapathy G, Yamazaki Y, Wolfstetter G, Mendoza P, Pfeifer K, Mohammed A, Hugosson F, Zhang H, Hsu AW et al (2015) FAM150A and FAM150B are activating ligands for anaplastic lymphoma kinase. Elife 4: e09811

Guan J, Yamazaki Y, Chand D, van Dijk JR, Ruuth K, Palmer RH, Hallberg B (2017) Novel Mechanisms of ALK Activation Revealed by Analysis of the Y1278S Neuroblastoma Mutation. Cancers (Basel) 9

Hafemeister C, Satija R (2019) Normalization and variance stabilization of single-cell RNA-seq data using regularized negative binomial regression. Genome Biol 20: 296

Hallberg B, Palmer RH (2013) Mechanistic insight into ALK receptor tyrosine kinase in human cancer biology. Nat Rev Cancer 13: 685–700

Hoehner JC, Gestblom C, Hedborg F, Sandstedt B, Olsen L, Pahlman S (1996) A developmental model of neuroblastoma: differentiating stroma-poor tumors’ progress along an extra-adrenal chromaffin lineage. Lab Invest 75: 659–675

Homem CCF, Steinmann V, Burkard TR, Jais A, Esterbauer H, Knoblich JA (2014) Ecdysone and mediator change energy metabolism to terminate proliferation in Drosophila neural stem cells. Cell 158: 874–888

Hu S, Fambrough D, Atashi JR, Goodman CS, Crews ST (1995) The Drosophila abrupt gene encodes a BTB-zinc finger regulatory protein that controls the specificity of neuromuscular connections. Genes Dev 9: 2936–2948

Huber K, Kalcheim C, Unsicker K (2009) The development of the chromaffin cell lineage from the neural crest. Auton Neurosci 151: 10–16

Ito K, Awano W, Suzuki K, Hiromi Y, Yamamoto D (1997) The Drosophila mushroom body is a quadruple structure of clonal units each of which contains a virtually identical set of neurones and glial cells. Development 124: 761–771

Ito K, Hotta Y (1992) Proliferation pattern of postembryonic neuroblasts in the brain of Drosophila melanogaster. Dev Biol 149: 134–148

Iwahara T, Fujimoto J, Wen D, Cupples R, Bucay N, Arakawa T, Mori S, Ratzkin B, Yamamoto T (1997) Molecular characterization of ALK, a receptor tyrosine kinase expressed specifically in the nervous system. Oncogene 14: 439–449

Janoueix-Lerosey I, Lequin D, Brugieres L, Ribeiro A, de Pontual L, Combaret V, Raynal V, Puisieux A, Schleiermacher G, Pierron G et al (2008) Somatic and germline activating mutations of the ALK kinase receptor in neuroblastoma. Nature 455: 967–970

Knoblich JA (2008) Mechanisms of asymmetric stem cell division. Cell 132: 583–597

Kuleshov MV, Jones MR, Rouillard AD, Fernandez NF, Duan Q, Wang Z, Koplev S, Jenkins SL, Jagodnik KM, Lachmann A et al (2016) Enrichr: a comprehensive gene set enrichment analysis web server 2016 update. Nucleic Acids Res 44: W90–97

Kumar A, Bello B, Reichert H (2009) Lineage-specific cell death in postembryonic brain development of Drosophila. Development 136: 3433–3442

Kumar S, Tunc I, Tansey TR, Pirooznia M, Harbison ST (2021) Identification of Genes Contributing to a Long Circadian Period in Drosophila Melanogaster. J Biol Rhythms 36: 239–253

Kurada P, White K (1998) Ras promotes cell survival in Drosophila by downregulating hid expression. Cell 95: 319–329

Lasek AW, Lim J, Kliethermes CL, Berger KH, Joslyn G, Brush G, Xue L, Robertson M, Moore MS, Vranizan K et al (2011) An evolutionary conserved role for anaplastic lymphoma kinase in behavioral responses to ethanol. PLoS One 6: e22636

Lee HH, Norris A, Weiss JB, Frasch M (2003) Jelly belly protein activates the receptor tyrosine kinase Alk to specify visceral muscle pioneers. Nature 425: 507–512

Lee T, Lee A, Luo L (1999) Development of the Drosophila mushroom bodies: sequential generation of three distinct types of neurons from a neuroblast. Development 126: 4065–4076

Lee T, Marticke S, Sung C, Robinow S, Luo L (2000) Cell-autonomous requirement of the USP/EcR-B ecdysone receptor for mushroom body neuronal remodeling in Drosophila. Neuron 28: 807–818

Liu LY, Long X, Yang CP, Miyares RL, Sugino K, Singer RH, Lee T (2019) Mamo decodes hierarchical temporal gradients into terminal neuronal fate. Elife 8

Liu Z, Yang CP, Sugino K, Fu CC, Liu LY, Yao X, Lee LP, Lee T (2015) Opposing intrinsic temporal gradients guide neural stem cell production of varied neuronal fates. Science 350: 317–320

Loren CE, Englund C, Grabbe C, Hallberg B, Hunter T, Palmer RH (2003) A crucial role for the Anaplastic lymphoma kinase receptor tyrosine kinase in gut development in Drosophila melanogaster. EMBO Rep 4: 781–786

Loren CE, Scully A, Grabbe C, Edeen PT, Thomas J, McKeown M, Hunter T, Palmer RH (2001) Identification and characterization of DAlk: a novel Drosophila melanogaster RTK which drives ERK activation in vivo. Genes Cells 6: 531–544

Madeira F, Park YM, Lee J, Buso N, Gur T, Madhusoodanan N, Basutkar P, Tivey ARN, Potter SC, Finn RD et al (2019) The EMBL-EBI search and sequence analysis tools APIs in 2019. Nucleic Acids Res 47: W636–W641

Maris JM (2010) Recent advances in neuroblastoma. N Engl J Med 362: 2202–2211

Marshall GM, Carter DR, Cheung BB, Liu T, Mateos MK, Meyerowitz JG, Weiss WA (2014) The prenatal origins of cancer. Nat Rev Cancer 14: 277–289

Martinsson T, Eriksson T, Abrahamsson J, Caren H, Hansson M, Kogner P, Kamaraj S, Schonherr C, Weinmar J, Ruuth K et al (2011) Appearance of the novel activating F1174S ALK mutation in neuroblastoma correlates with aggressive tumor progression and unresponsiveness to therapy. Cancer Res 71: 98–105

Mendoza-Garcia P, Hugosson F, Fallah M, Higgins ML, Iwasaki Y, Pfeifer K, Wolfstetter G, Varshney G, Popichenko D, Gergen JP et al (2017) The Zic family homologue Odd-paired regulates Alk expression in Drosophila. PLoS Genet 13: e1006617

Michki NS, Li Y, Sanjasaz K, Zhao Y, Shen FY, Walker LA, Cao W, Lee CY, Cai D (2021) The molecular landscape of neural differentiation in the developing Drosophila brain revealed by targeted scRNA-seq and multi-informatic analysis. Cell Rep 35: 109039

Mo ES, Cheng Q, Reshetnyak AV, Schlessinger J, Nicoli S (2017a) Alk and Ltk ligands are essential for iridophore development in zebrafish mediated by the receptor tyrosine kinase Ltk. Proc Natl Acad Sci U S A 114: 12027–12032

Mo ES, Cheng QN, Reshetnyak AV, Schlessinger J, Nicoli S (2017b) Alk and Ltk ligands are essential for iridophore development in zebrafish mediated by the receptor tyrosine kinase Ltk. P Natl Acad Sci USA 114: 12027–12032

Mosse YP, Laudenslager M, Longo L, Cole KA, Wood A, Attiyeh EF, Laquaglia MJ, Sennett R, Lynch JE, Perri P et al (2008) Identification of ALK as a major familial neuroblastoma predisposition gene. Nature 455: 930–935

Muller HA (2008) Immunolabeling of embryos. Methods Mol Biol 420: 207–218

Okamoto N, Nishimura T (2015) Signaling from Glia and Cholinergic Neurons Controls Nutrient-Dependent Production of an Insulin-like Peptide for Drosophila Body Growth. Dev Cell 35: 295–310

Ono S, Saito T, Terui K, Yoshida H, Enomoto H (2019) Generation of conditional ALK F1174L mutant mouse models for the study of neuroblastoma pathogenesis. Genesis 57: e23323

Orthofer M, Valsesia A, Magi R, Wang QP, Kaczanowska J, Kozieradzki I, Leopoldi A, Cikes D, Zopf LM, Tretiakov EO et al (2020) Identification of ALK in Thinness. Cell 181: 1246–1262 e1222

Ouyang JF, Kamaraj US, Cao EY, Rackham OJL (2021) ShinyCell: Simple and sharable visualisation of single-cell gene expression data. Bioinformatics (Oxford, England)

Öztürk-Çolak A, Moussian B, Araújo SJ, Casanova J (2016) A feedback mechanism converts individual cell features into a supracellular ECM structure in Drosophila trachea. eLife 5: e09373

Paskus JD, Herring BE, Roche KW (2020) Kalirin and Trio: RhoGEFs in Synaptic Transmission, Plasticity, and Complex Brain Disorders. Trends Neurosci 43: 505–518

Pecot MY, Chen Y, Akin O, Chen Z, Tsui CY, Zipursky SL (2014) Sequential axon-derived signals couple target survival and layer specificity in the Drosophila visual system. Neuron 82: 320–333

Pignoni F, Zipursky SL (1997) Induction of Drosophila eye development by decapentaplegic. Development 124: 271–278

Pinto-Teixeira F, Konstantinides N, Desplan C (2016) Programmed cell death acts at different stages of Drosophila neurodevelopment to shape the central nervous system. FEBS Lett 590: 2435–2453

Ren Q, Yang CP, Liu Z, Sugino K, Mok K, He Y, Ito M, Nern A, Otsuna H, Lee T (2017) Stem Cell-Intrinsic, Seven-up-Triggered Temporal Factor Gradients Diversify Intermediate Neural Progenitors. Curr Biol 27: 1303–1313

Reshetnyak AV, Murray PB, Shi X, Mo ES, Mohanty J, Tome F, Bai H, Gunel M, Lax I, Schlessinger J (2015) Augmentor alpha and beta (FAM150) are ligands of the receptor tyrosine kinases ALK and LTK: Hierarchy and specificity of ligand-receptor interactions. Proc Natl Acad Sci U S A 112: 15862–15867

Rohrbough J, Broadie K (2010) Anterograde Jelly belly ligand to Alk receptor signaling at developing synapses is regulated by Mind the gap. Development 137: 3523–3533

Rohrbough J, Kent KS, Broadie K, Weiss JB (2013) Jelly Belly trans-synaptic signaling to anaplastic lymphoma kinase regulates neurotransmission strength and synapse architecture. Dev Neurobiol 73: 189–208

Rossi AM, Desplan C (2020) Extrinsic activin signaling cooperates with an intrinsic temporal program to increase mushroom body neuronal diversity. Elife 9

Rossini L, Contarini M, Giarruzzo F, Assennato M, Speranza S (2020) Modelling Drosophila suzukii Adult Male Populations: A Physiologically Based Approach with Validation. Insects 11

Saito D, Takase Y, Murai H, Takahashi Y (2012) The dorsal aorta initiates a molecular cascade that instructs sympatho-adrenal specification. Science 336: 1578–1581

Stuart T, Butler A, Hoffman P, Hafemeister C, Papalexi E, Mauck WM, 3rd, Hao Y, Stoeckius M, Smibert P, Satija R (2019) Comprehensive Integration of Single-Cell Data. Cell 177: 1888–1902 e1821

Stute C, Schimmelpfeng K, Renkawitz-Pohl R, Palmer RH, Holz A (2004) Myoblast determination in the somatic and visceral mesoderm depends on Notch signalling as well as on milliways(mili(Alk)) as receptor for Jeb signalling. Development 131: 743–754

Syed MH, Mark B, Doe CQ (2017a) Playing Well with Others: Extrinsic Cues Regulate Neural Progenitor Temporal Identity to Generate Neuronal Diversity. Trends Genet 33: 933–942

Syed MH, Mark B, Doe CQ (2017b) Steroid hormone induction of temporal gene expression in Drosophila brain neuroblasts generates neuronal and glial diversity. Elife 6

Tolbert VP, Matthay KK (2018) Neuroblastoma: clinical and biological approach to risk stratification and treatment. Cell Tissue Res 372: 195–209

Tomolonis JA, Agarwal S, Shohet JM (2018) Neuroblastoma pathogenesis: deregulation of embryonic neural crest development. Cell Tissue Res 372: 245–262

Trigg RM, Turner SD (2018) ALK in Neuroblastoma: Biological and Therapeutic Implications. Cancers (Basel) 10

Umapathy G, Mendoza-Garcia P, Hallberg B, Palmer RH (2019) Targeting anaplastic lymphoma kinase in neuroblastoma. APMIS

Urbach R, Technau GM (2004) Neuroblast formation and patterning during early brain development in Drosophila. Bioessays 26: 739–751

Varshney GK, Palmer RH (2006) The bHLH transcription factor Hand is regulated by Alk in the Drosophila embryonic gut. Biochem Biophys Res Commun 351: 839–846

Vernersson E, Khoo NK, Henriksson ML, Roos G, Palmer RH, Hallberg B (2006) Characterization of the expression of the ALK receptor tyrosine kinase in mice. Gene Expr Patterns 6: 448–461

Vieceli FM, Bronner ME (2018) Leukocyte receptor tyrosine kinase interacts with secreted midkine to promote survival of migrating neural crest cells. Development 145

Weiss JB, Suyama KL, Lee HH, Scott MP (2001) Jelly belly: a Drosophila LDL receptor repeat-containing signal required for mesoderm migration and differentiation. Cell 107: 387–398

Weiss JB, Weber S, Marzulla T, Raber J (2017) Pharmacological inhibition of Anaplastic Lymphoma Kinase rescues spatial memory impairments in Neurofibromatosis 1 mutant mice. Behav Brain Res 332: 337–342

Weiss JB, Xue C, Benice T, Xue L, Morris SW, Raber J (2012) Anaplastic lymphoma kinase and leukocyte tyrosine kinase: functions and genetic interactions in learning, memory and adult neurogenesis. Pharmacol Biochem Behav 100: 566–574

Weng M, Golden KL, Lee CY (2010) dFezf/Earmuff maintains the restricted developmental potential of intermediate neural progenitors in Drosophila. Dev Cell 18: 126–135

Witek B, El Wakil A, Nord C, Ahlgren U, Eriksson M, Vernersson-Lindahl E, Helland A, Alexeyev OA, Hallberg B, Palmer RH (2015) Targeted Disruption of ALK Reveals a Potential Role in Hypogonadotropic Hypogonadism. PLoS One 10: e0123542

Wolf FA, Angerer P, Theis FJ (2018) SCANPY: large-scale single-cell gene expression data analysis. Genome Biol 19: 15

Wolfstetter G, Pfeifer K, Backman M, Masudi TA, Mendoza-García P, Chen S, Sonnenberg H, Sukumar SK, Uçkun E, Varshney GK et al (2020) Identification of the Wallenda JNKKK as an Alk suppressor reveals increased competitiveness of Alk-expressing cells. Scientific Reports 10: 14954

Wolfstetter G, Pfeifer K, van Dijk JR, Hugosson F, Lu X, Palmer RH (2017) The scaffolding protein Cnk binds to the receptor tyrosine kinase Alk to promote visceral founder cell specification in Drosophila. Sci Signal 10

Woodling NS, Aleyakpo B, Dyson MC, Minkley LJ, Rajasingam A, Dobson AJ, Leung KHC, Pomposova S, Fuentealba M, Alic N et al (2020) The neuronal receptor tyrosine kinase Alk is a target for longevity. Aging Cell 19: e13137

Yang CP, Fu CC, Sugino K, Liu Z, Ren Q, Liu LY, Yao X, Lee LP, Lee T (2016) Transcriptomes of lineage-specific Drosophila neuroblasts profiled by genetic targeting and robotic sorting. Development 143: 411–421

Yang CP, Samuels TJ, Huang Y, Yang L, Ish-Horowicz D, Davis I, Lee T (2017) Imp and Syp RNA-binding proteins govern decommissioning of Drosophila neural stem cells. Development 144: 3454–3464

Yao S, Cheng M, Zhang Q, Wasik M, Kelsh R, Winkler C (2013) Anaplastic lymphoma kinase is required for neurogenesis in the developing central nervous system of zebrafish. PLoS One 8: e63757

Yu F, Kuo CT, Jan YN (2006) Drosophila neuroblast asymmetric cell division: recent advances and implications for stem cell biology. Neuron 51: 13–20

Zhu S, Lee JS, Guo F, Shin J, Perez-Atayde AR, Kutok JL, Rodig SJ, Neuberg DS, Helman D, Feng H et al (2012) Activated ALK collaborates with MYCN in neuroblastoma pathogenesis. Cancer Cell 21: 362–373

Zielke N, Korzelius J, van Straaten M, Bender K, Schuhknecht GFP, Dutta D, Xiang J, Edgar BA (2014) Fly-FUCCI: A versatile tool for studying cell proliferation in complex tissues. Cell Rep 7: 588–598

Zong W, Liu S, Wang X, Zhang J, Zhang T, Liu Z, Wang D, Zhang A, Zhu M, Gao J (2015) Trio gene is required for mouse learning ability. Brain Res 1608: 82–90

